# Understanding the biosynthesis, metabolic regulation, and anti-phytopathogen activity of 3,7-dihydroxytropolone in *Pseudomonas* spp

**DOI:** 10.1101/2024.04.03.587903

**Authors:** Alaster D. Moffat, Lars Höing, Javier Santos-Aberturas, Tim Markwalder, Jacob G. Malone, Robin Teufel, Andrew W. Truman

## Abstract

The genus *Pseudomonas* is a prolific source of specialized metabolites with significant biological activities, including siderophores, antibiotics, and plant hormones. These molecules play pivotal roles in environmental interactions, influencing pathogenicity, inhibiting microorganisms, responding to nutrient limitation and abiotic challenges, and regulating plant growth. These properties mean that pseudomonads are candidates as biological control agents against plant pathogens. Multiple transposon-based screens have identified a *Pseudomonas* biosynthetic gene cluster (BGC) associated with potent antibacterial and antifungal activity that produces 7-hydroxytropolone (7-HT). In this study, we show that this BGC also makes 3,7-dihydroxytropolone (3,7-dHT), which has strong antimicrobial activity towards *Streptomyces scabies*, a potato pathogen. Both molecules exhibit broad biological activities, suggesting roles in competitive soil and plant microbial communities. Through metabolomics and reporter assays, we unveil the involvement of cluster-situated genes in generating phenylacetyl-coenzyme A, a key precursor for tropolone biosynthesis via the phenylacetic acid catabolon. The clustering of these phenylacetic acid genes within tropolone BGCs is unusual in other Gram-negative bacteria. Our findings support the interception of phenylacetic acid catabolism via an enoyl-CoA dehydratase encoded in the BGC, as well as highlighting an essential biosynthetic role for a conserved thioesterase. Biochemical assays were used to show that this thioesterase functions after a dehydrogenation-epoxidation step catalysed by a flavoprotein. We use this information to identify diverse uncharacterised BGCs that encode proteins with homology to flavoproteins and thioesterases involved in tropolone biosynthesis. This study provides insights into tropolone biosynthesis in *Pseudomonas*, laying the foundation for further investigations into the ecological role of tropolone production.

## INTRODUCTION

The bacterial genus *Pseudomonas* is widely distributed in nature, where it is ubiquitously found in soils worldwide^1^, with extraordinary inter-species and intra-species diversity that is reflected in a pangenome of over 60,000 unique genes^2,3^. As a result of this tremendous diversity, *Pseudomonas* spp. are a well-known source of novel specialised metabolites (SMs) with important biological activities^4–6^, including siderophores like pyoverdine and pyochelin^7^, antibiotics such as obafluorin^8^ and mupirocin^9^, extracellular electron shuttles such as phenazines^10^, as well as a diverse repertoire of volatile compounds^11^. *Pseudomonas* SMs often have critical roles in mediating interactions with other organisms in the environment^12,13^, including toxins associated with pathogenicity, antibiotics and antifungals to inhibit other microorganisms, and hormones that influence plant growth and development^12^. The capacity of *Pseudomonas* spp. to colonise plant roots and inhibit other organisms means that many strains of this genus have already been targeted for use in crop protection programmes as biological control agents that can suppress plant pathogens^14^.

A significant number of SM biosynthetic gene clusters (BGCs) do not possess canonical features that enable detection by widely used genome mining tools and are instead discovered using activity-guided transposon mutagenesis screens. Recently, several independent high-throughput transposon-based screens discovered a *Pseudomonas* BGC associated with potent antibacterial and antifungal activity, including our own work showing its involvement in the inhibition of *Streptomyces scabies*, a bacterium that causes potato scab, by *Pseudomonas* sp. Ps652^15^. This BGC was initially identified by Xie and co-workers using a transposon screen of *Pseudomonas donghuensis* HYS to identify the BGC of a non-fluorescent siderophore^16^, which they later determined to be 7-hydroxytropolone^17,18^ (7-HT). This tropolone BGC was also independently identified by transposon mutagenesis as the key determinant of suppressive activity of *P. donghuensis* isolates towards the bacterium *Pectobacterium carotovorum*^19^ (potato black leg and soft rot) and the fungus *Macrophomina phaseolina*^20^ (broad spectrum stem and root rot), as well as the fungus *Verticillium dahliae* (cotton verticillium wilt) via targeted inactivation of the BGC^21^.

Tropolones are small molecules containing an aromatic 7-membered cyclohepta-2,4,6-trienone (tropone) ring that is hydroxylated at position 2 (Figure 1). Tropolone SMs have been documented in plants, fungi, and bacteria, and possess an extensive range of biological activities, including antibacterial, antifungal, antiviral and cytotoxic activities^22,23^. In bacteria, tropolone biosynthesis has been a long-standing question since the first discovery of tropolone production by *Pseudomonas* ATCC 31099 in 1980^24^. Some larger tropolone-containing SMs like the isatropolones (Figure 1) are known to be produced via a polyketide synthase (PKS) route^25^, while genetic evidence for the route to the standalone tropolone skeleton was first determined via transposon mutagenesis to identify genes in the Rhodobacteraceae involved in the biosynthesis of tropodithietic acid, a sulfur-containing tropone^26^.

**Figure 1.**
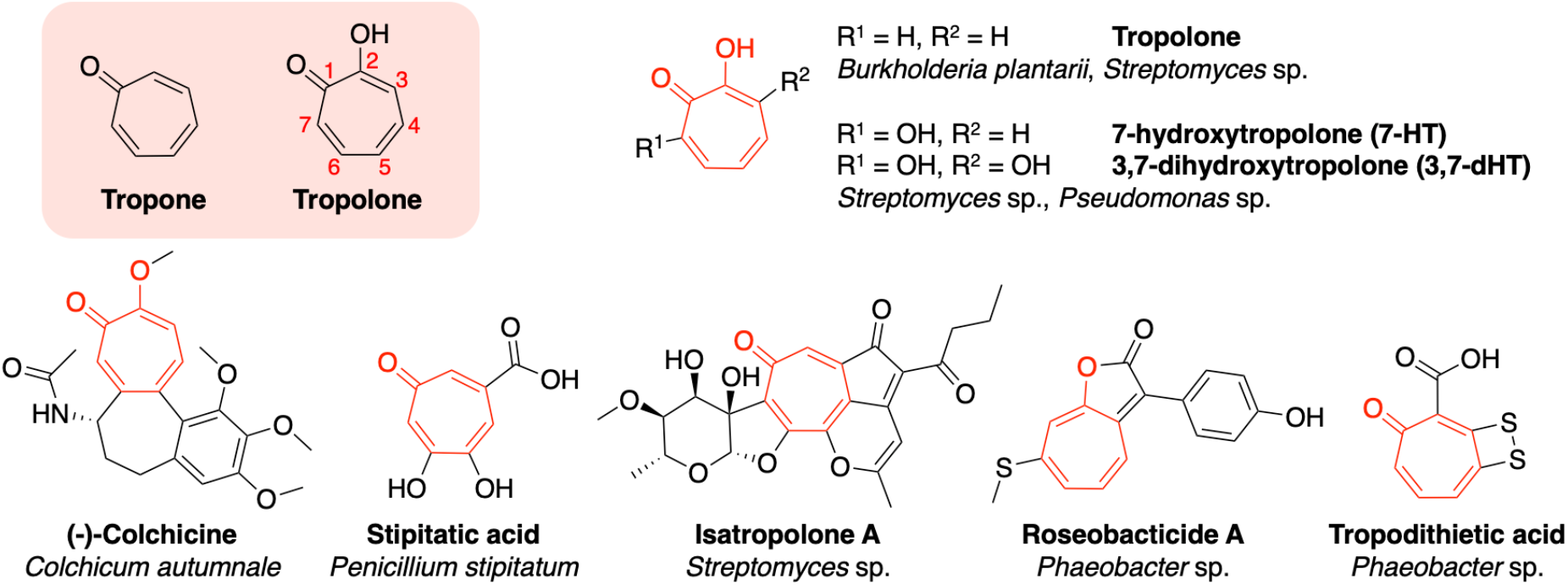
The tropone and tropolone moieties alongside examples of tropolone natural products and their diverse bacterial, fungal and plant producers.

Further studies demonstrated that tropodithietic acid derives from phenylacetic acid (PAA) catabolism^27,28^ via a key branching point^29–32^ to direct flux of primary metabolic intermediates towards the biosynthesis of a 7-membered ring tropolone precursor, which was first proposed in 1992 following isotope feeding experiments in thiotropocin biosynthesis^33^. A conceptually similar pathway occurs in *Streptomyces*, which also branches from the PAA degradative pathway, followed by additional modifications by dedicated biosynthetic enzymes, but this BGC appears to be distinct to the Gram-negative tropolone BGCs found in *Phaeobacter*, *Burkholderia* and *Pseudomonas*. We recently described a novel class of dioxygenase in *Burkholderia* capable of producing tropolone via the PAA catabolon^34^, but questions still remain about earlier biosynthetic steps, the full complement of genes required for the production of tropolones, and the function of enzymes catalysing later stage modifications, such as additional hydroxylation or incorporation of sulphur atoms. In particular, very little is known about tropolone biosynthesis in *Pseudomonas*.

In this study we report that the tropolone (*tpo*) BGC in *Pseudomonas* sp. Ps652 is responsible for the production of 3,7-dihydroxytropolone (3,7-dHT), which may represent the true product of the *tpo* BGC. This is the first time the production of 3,7-dHT has been reported for a Gram-negative bacterium. This metabolite is a critical determinant of suppressive activity towards the potato pathogens *S. scabies*, a bacterium that causes common scab, and *Phytophthora infestans*, an oomycete that causes potato late blight. We show that 3,7-dHT has greater potency against *S. scabies* 87-22 when compared to 7-HT and investigate the role of a conserved thiamine pyrophosphate (TPP) - dependent decarboxylase and a phenylacetyl-coenzyme A (PA-CoA) ligase in the early stages of biosynthesis and regulation of the pathway. These two genes form part of the core BGC, serving to derepress the PAA catabolon under conditions of low environmental PAA availability, thereby allowing production of tropolones in those conditions. We also demonstrate that a conserved thioesterase has a key role alongside a flavoprotein for a late-stage step in tropolone biosynthesis and use this information to identify new BGCs.

## RESULTS AND DISCUSSION

### Gene deletions highlight the importance of tropolone production for suppressive activity

Our prior transposon mutagenesis study^15^ highlighted an important role for the *tpo* BGC in the inhibition of *S. scabies* by *Pseudomonas* sp. Ps652 (herein simply referred to as Ps652) an environmental isolate from a commercial potato field in Norfolk (U.K.). However, that transposon screen may have overlooked the contributions of the other BGCs, especially towards other plant pathogens. To ensure that BGCs were accurately annotated, we obtained a higher quality single-scaffold genome assembly, produced by Illumina and Oxford Nanopore sequencing (accession OZ024668.1). Multi-locus sequence typing (MLST) using AutoMLST^35^ (Figure S1) showed that Ps652 possesses 93.5% average nucleotide identity (ANI) to *Pseudomonas* sp. UC 17F4 (GCF_900101695) and 93.4% to *Pseudomonas donghuensis* (GCF_000259195), a previously characterised tropolone producer^18^. These ANIs fall below the 94.0% threshold set for the distinction of species in *Pseudomonas*^36,37^. Accordingly, it is likely that Ps652 represents a novel species of *Pseudomonas*, which is also supported by an analysis using the Type Strain Genome Server^38^. Phylogenetic analysis places it within the *P. vranovensis* sub-group within the *P. putida* group^39^ (Figure S1).

The BGC detection tool antiSMASH 7.0.0^40^ detected 10 BGCs in the Ps652 genome (Figure 2A, Table S5), including BGCs predicted to be involved in the biosynthesis of the siderophore pyoverdine, an aryl polyene^41^, the redox cofactor pyrroloquinoline quinone^42^ (PQQ), and a homologue of the *Pseudomonas virulence factor* (*pvf*) BGC, which is predicted to make autoinducers that regulate gene expression^43^. Analysis using GECCO, an alternative BGC identification tool^44^, revealed two more putative BGCs (Table S5) but neither tool detected the tropolone BGC. As reported previously^13^, further analysis of the genome revealed genes for the biosynthesis of hydrogen cyanide^45^ (HCN), hydrogen sulfide^46^ and indole-3-acetic acid^47^ (IAA), a common plant auxin (Figure 2A).

**Figure 2.**
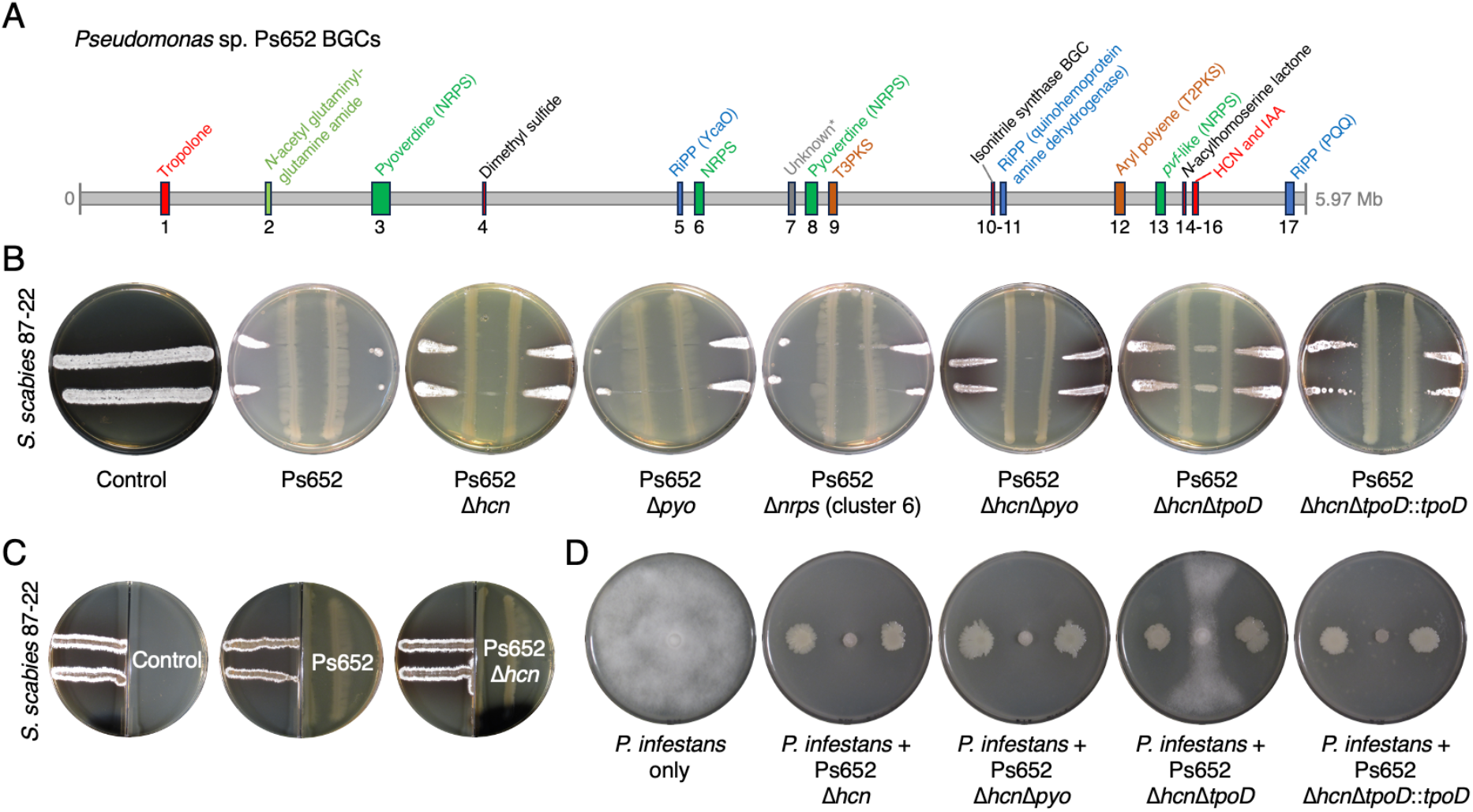
Ps652 BGCs and the effect of deletions on suppressive activity. A. Overview of BGCs identified from antiSMASH 7.0, GECCO 0.9.8 and manual annotation. *Unknown relates to a GECCO hit with a low number of biosynthetic genes. Table S5 provides further BGC details. B. Cross-streak assays between Ps652 mutants and *S. scabies*. C. Split-plate assays assessing the role of volatiles in *S. scabies* inhibition. No change in *S. scabies* morphology (left hand side) was detected with either Ps652 or the *hcn* deletion mutant. D. Assays between *P. infestans* (central growth) and Ps652 mutants. Further split-plate assays are shown in Figure S2.

Deletion of the HCN BGC had a negligible effect on biological activity towards *S. scabies* (Figure 2B) and *P. infestans* (Figure 2D), while in-frame deletions in the pyoverdine BGC (Δ*pyo*) and a BGC encoding a standalone NRPS module (Δ*nrps*, BGC6, Figure 2A, Table S5) indicated that these BGCs were not directly involved in suppressive activity. Split plate assays using plates with an internal barrier (Figure 2C) indicated that volatiles were not major determinants of suppression towards either pathogen, although volatile activity was medium dependent for *P. infestans* inhibition (Figure S2), highlighting the production of further suppressive molecules.

A gene naming scheme for the *Pseudomonas* tropolone BGC (*tpo*) is proposed based on BGC boundaries inferred from mutants generated in this study and prior studies on homologous BGCs in other *Pseudomonas* strains^15,16,18,20,21^ (Figure 3A, Table S6). To support the role of the *tpo* BGC in *S. scabies* inhibition and to rule out the possibility that prior transposon mutants had unintended polar effects on Ps652, an in-frame deletion of the *tpoD* gene was generated. All mutants were generated in Ps652 Δ*hcn* to eliminate the possibility that HCN-based inhibition obscured *tpo*-dependent effects in subsequent assays. *tpoD* encodes a predicted acyl-CoA thioesterase and homologues are conserved across previously identified tropolone BGCs in Gram-negative bacteria but without a documented role to date. This Δ*hcn*Δ*tpoD* mutant resulted in the loss of suppression of *S. scabies* 87-22 growth at a distance, where *S. scabies* growth could be observed between the Ps652 streaks (Figure 2B) and could be genetically complemented by expression of *tpoD* from plasmid pME6032^48^ (Figure 2B). Suppressive activity towards *P. infestans* was also significantly reduced in Ps652 Δ*hcn*Δ*tpoD*, although a substantial clearance zone remained, indicating that there are further anti-oomycete SMs produced by Ps652 (Figure 2D).

**Figure 3.**
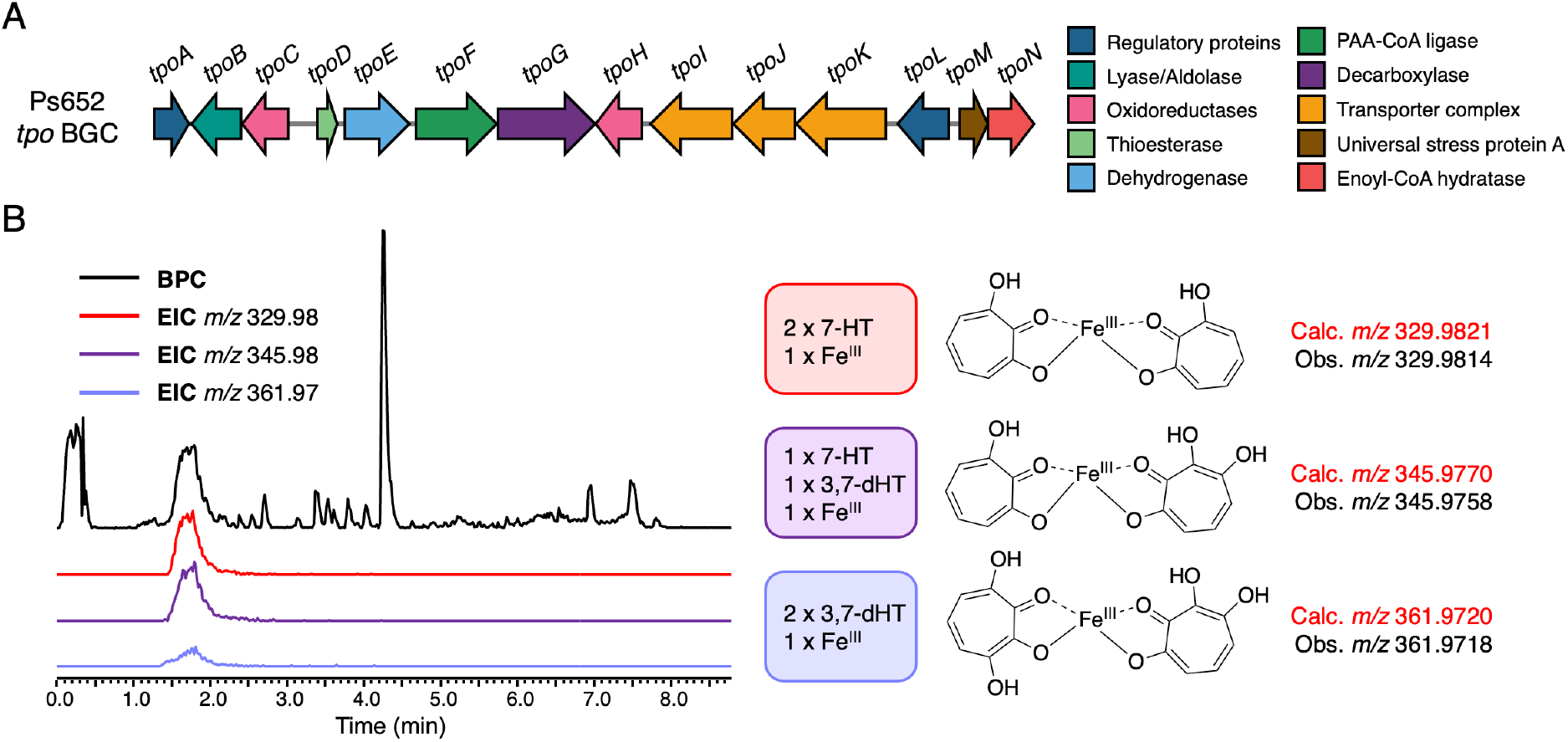
Tropolone biosynthesis in Ps652. A. The tropolone BGC in *Pseudomonas* sp. Ps652 and a new suggested naming scheme. B. Tropolone-iron chelates observed in LC-MS data of extracts from Ps652 Δ*hcn* that suggested the production of an additional dihydroxylated tropolone (BPC = base peak chromatogram; EIC = extracted ion chromatogram).

### *Pseudomonas* sp. Ps652 produces multiple tropolones

After discovery of the tropolone BGC in Ps652, and confirmation of its link to suppressive activity towards *S. scabies* 87-22 and *P. infestans*, it was essential to establish whether Ps652 indeed produces tropolones. Using a synthetic standard of 7-HT for comparison in liquid chromatography – mass spectrometry (LC-MS) analyses, we were able to detect 7-HT (calculated [M+H]^+^ *m/z* 139.0390; observed *m/z* 139.0389) in extracts of Ps652 (Figure S3). 7-HT was undetectable in the equivalent extracts from Ps652 Δ*hcn*Δ*tpoD*, confirming its identity as a hydroxylated tropolone related to the BGC. However, we also observed some molecules with the characteristic 330 nm absorbance of tropolones but with larger mass-to-charge ratios (Figure 3B, Figure S3) that were consistent with the masses of tropolone-iron chelates, matching the 2:1 tropolone to iron stoichiometry demonstrated by Jiang *et al*.^17^ Interestingly, some of these masses corresponded to the predicted masses of a dihydroxylated tropolone in complex with iron and 7-HT (calculated *m/z* 345.9770; observed *m/z* 345.9760), or two molecules of a dihydroxylated tropolone in complex with iron(III) (calculated *m/z* 361.9718; observed *m/z* 361.9719) (Figure S3). The corresponding monomer of the dihydroxylated tropolone was subsequently detected in MS data ([M+H]^+^, calculated *m/z* 155.0339; observed *m/z* 155.0339) (Figure 4A, Figure S4).

**Figure 4.**
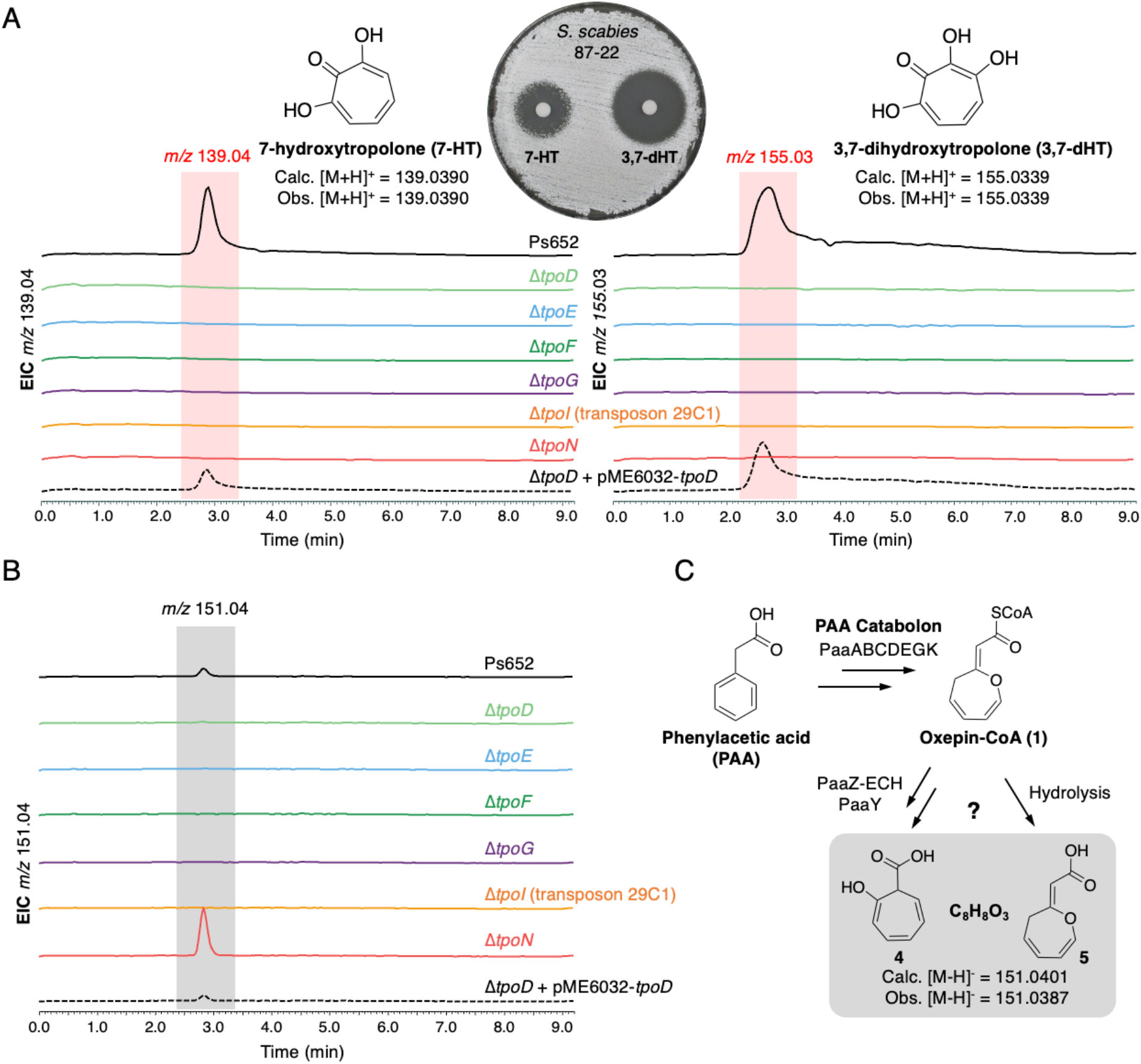
Gene deletions in the Ps652 tropolone BGC. A. LC-MS spectra showing production of 7-HT and 3,7-HT by Ps652. Deletion of *tpoD* abolishes production of tropolones, which can be restored by genetic complementation with pME6032-*tpoD*. Production is also abolished across each tropolone BGC mutant. Each mutant is generated in Ps652 Δ*hcn*. 3,7-dHT is more active against *S. scabies* compared to 7-HT (10 μg each compound used in assay). B. Production of m/z 151.04 molecule (negative mode MS) across BGC mutants. C. Routes to two possible shunt metabolites that have a mass consistent with the observed molecule.

We purified this dihydroxylated tropolone from Ps652 Δ*hcn*Δ*pyo* grown in 2 litres of liquid modified King’s Broth plus glucose (MKBG, Table S4) using size-exclusion chromatography and high-performance liquid chromatography (HPLC, Figure S4), to yield 2.8 mg of pure material. NMR analysis (Figures S5-S6) identified the molecule as 3,7-dihydroxytropolone (3,7-dHT), matching a previously published characterisation^49^ with ∂_C_ = 119.2 ppm, 129.4 ppm, 158.1 ppm and 158.9 ppm. 3,7-dHT has an ability to bind iron equivalent to or greater than 7-HT in a qualitative assay on chrome azurol S (CAS) agar^50^ (Figure S7), suggesting it is also capable of functioning as a siderophore like 7-HT^17^. Additionally, disk diffusion assays against *S. scabies* 87-22 indicated that 3,7-dHT is a more potent antibiotic compared to 7-HT (Figure 4A), with respective minimum inhibitory concentrations (MICs) of 2.5 and 5 µg/mL (Figure S7). Together, these data suggest that 3,7-dHT may constitute the true final product of the tropolone BGC in *Pseudomonas*, whereas 7-HT could be a biosynthetic intermediate or shunt metabolite lacking the final hydroxylation. No additionally hydroxylated tropolones were observed in our LC-MS data.

### Understanding tropolone biosynthesis in *Pseudomonas*

In order to understand which genes are genuinely required for tropolone biosynthesis, we generated a series of nonpolar deletion mutants to supplement previously generated transposon mutants, which included *tpoI* (encoding a putative transporter component; transposon mutant 29C1). In addition to *tpoD* (encoding a putative thioesterase), in-frame mutants were also generated in Ps652 Δ*hcn* for genes encoding TpoE (dehydrogenase-like flavoprotein), TpoF (acyl-CoA ligase), TpoG (TPP-dependent decarboxylase) and TpoN (enoyl-CoA dehydratase). All mutants were unable to make 7-HT and 3,7-dHT when grown in MKBG, demonstrating the involvement of the deleted genes in tropolone biosynthesis (Figure 4A). Genetic complementation of *tpoE* and *tpoF* using pUC18-mini-Tn7-based plasmids^51^ restored tropolone biosynthesis (Figure S8), indicating that there were no unanticipated secondary effects of deleting these mid-operon genes.

However, we were unable to identify intermediates or shunt metabolites from most mutants via targeted and untargeted analyses of LC-MS spectra. The lack of detectable molecules could be caused by compound instability, volatility, and/or the disruption of early biosynthetic steps. One exception was Δ*tpoN*, where a molecule of *m/z* 151.04 was observed in negative mode MS. This compound was detected in extracts of parental Ps652 Δ*hcn*, but in substantially lower amounts than in Δ*tpoN* (Figure 4B). TpoN is an enoyl-CoA hydratase (ECH), which is one of the domains of PaaZ, a dual domain protein involved in PAA catabolism that features an ECH domain and an aldehyde dehydrogenase (ALDH) domain. PaaZ catalyses the ring opening and oxidation of (*Z*)-2-(oxepin-2(3*H*)-ylidene)-acetyl CoA (oxepin-CoA, **1**), a key intermediate in the PAA catabolon. In the absence of a functional ALDH domain, the PaaZ ECH domain catalyses the formation of a highly reactive 3-oxo-5,6-dehydrosuberoyl-CoA semialdehyde (**2**) that can undergo a spontaneous Knoevenagel condensation to generate a 7-membered tropolone precursor (2-hydroxycyclohepta-1,4,6-triene-1-formyl-CoA, **3**)^29^ ^52^ (Figure 5).

**Figure 5.**
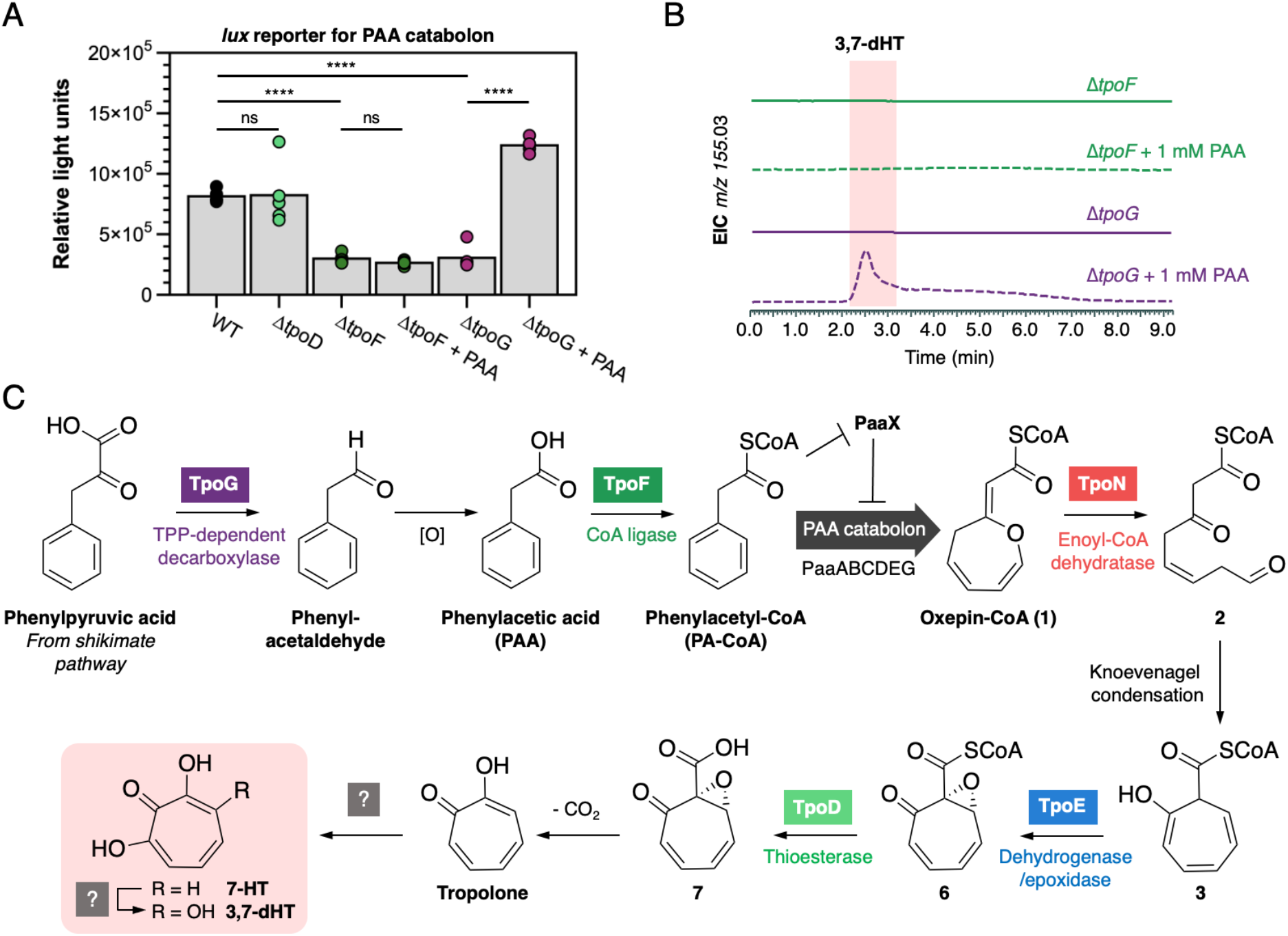
Role of TpoF and TpoG in tropolone biosynthesis. A. Luciferase (*lux*) reporter assay for the expression of the PAA catabolon. The parental “WT” strain is Ps652 Δ*hcn*. PAA was added at a concentration of 1 mM. Data were collected for 5 replicates for each condition, which are presented along with the respective means (grey bars). Significance determined by Student’s *t*-test (two-tailed, unequal variance) is presented: ns = not significant (*p*>0.05), **** = *p*<0.0001. B. LC-MS spectra showing chemical complementation of Δ*tpoG* with exogenous PAA, whereas production is not restored when PAA is fed to Δ*tpoF*. C. Proposed biosynthetic scheme for 7-HT and 3,7-dHT in Ps652.

The instability of the *m/z* 151.04 molecule prevented full characterisation, but the accurate mass is consistent with 2-hydroxycyclohepta-1,4,6-triene-1-carboxylic acid (**4**) (C_8_H_8_O_3_, calculated [M-H]^−^ *m/z* 151.0401; observed *m/z* 151.0387), which is a shunt metabolite that can be formed by the PAA catabolon via the thioesterase PaaY (Figure 4C). Alternatively, thioester hydrolysis of **1** would also provide a C_8_H_8_O_3_ molecule (**5**), which could occur if the PAA catabolon stalls prior to PaaZ/TpoN activity (Figure 4C). In *Streptomyces cyaneofuscatus*, a standalone ECH is proposed to direct PAA flux towards tropolone biosynthesis^53^, and our data supports an equivalent function for TpoN in *Pseudomonas* given the production of a putative shunt metabolite from the PAA catabolon.

### Role of phenylacetate-CoA ligase (TpoF) and a TPP-dependent decarboxylase (TpoG) in tropolone biosynthesis in *Pseudomonas*

Gene deletions in Ps652 highlighted an essential role for *tpoF* and *tpoG* in tropolone biosynthesis. Prior transposon mutagenesis and gene regulation studies have reported *tpoF* and *tpoG* homologues as important for biological activity and siderophore phenotypes related to tropolone biosynthesis in various *Pseudomonas* strains^16,18,20^. However, to date, no precise biosynthetic roles have been proposed and homologues of these genes are not found in tropolone BGCs in other Gram-negative bacteria. TpoF is a paralog of the phenylacetyl-CoA (PA-CoA) ligase PaaK from the PAA catabolon. It has been shown in *Escherichia coli* that *paaK* and the rest of the PAA catabolon is only expressed in the presence of PA-CoA, which directly depresses PaaX, a transcriptional repressor that controls the PAA catabolon^52^. TpoG is a putative TPP-dependent decarboxylase (Table S6), which is the same protein family (COG3961) as IpdC from *Azospirillum brasilense*. IpdC catalyses the decarboxylation of phenylpyruvate to phenylacetaldehyde as part of auxin biosynthesis^54,55^. Phenylacetaldehyde could then be further oxidised to PAA, potentially via a non-clustered phenylacetaldehyde dehydrogenase (Ps652 VVM71318.1 is 52% identical to phenylacetaldehyde dehydrogenase from the *Pseudomonas putida* S12 styrene catabolic pathway^56^). We therefore hypothesised that, in the absence of PAA or PA-CoA from primary metabolism, TpoG and TpoF would cooperate to supply PA-CoA as a tropolone precursor and to activate expression of the PAA catabolon, which is required for tropolone production.

To test this hypothesis, a PAA catabolon reporter was generated by fusing the promoter of *paaA* from Ps652 to the luciferase (*lux*) operon, which generates bioluminescence when expressed ^57^. Use of this *paa* promoter-*lux* fusion reporter in Δ*tpoF* and Δ*tpoG* indicated that expression of the PAA catabolon was significantly lower than in the parental Ps652 Δ*hcn* strain (Figure 5A). Additionally, we observed that supplying exogenous PAA was not sufficient to chemically complement the Δ*tpoF* mutant but PAA could complement the Δ*tpoG* mutant (Figure 5B). These results are consistent with TpoG directly increasing the production of PAA and TpoF then converting PAA to PA-CoA. PA-CoA could then function to derepress the PAA catabolon by rendering PaaX inactive. When exogenous PAA is added, PA-CoA can be produced by TpoF in the Δ*tpoG* mutant but not in the Δ*tpoF* mutant. TpoF and TpoG thus function to supply an essential precursor and activate the PAA catabolon for tropolone production in the absence of environmental PAA (Figure 5C). A similar situation is seen in the biosynthesis of tropolones in *S. cyaneofuscatus*, where TrlB and TrlH help shuttle primary metabolites towards the PAA catabolon and ultimately tropolone biosynthesis^53^, although these proteins have different catalytic roles to TpoF and TpoG.

### TpoD and TpoE function to generate an advanced tropolone precursor

TpoD (thioesterase) and TpoE (flavoprotein) encoded in the *Pseudomonas tpo* BGC are essential for tropolone biosynthesis (Figure 4A). TpoE is a homologue of flavoproteins previously shown to be critical for tropodithietic acid biosynthesis in *Phaeobacter inhibens* (TdaE^Pi^) and tropolone biosynthesis in *Burkholderia plantarii* (TdaE^Bp^)^34^. TdaE^Pi^ and TdaE^Bp^ homologues were furthermore shown to convert **3** into (2*R*,3*R*)-epoxytropone-2-carboxylate (**7**, Figure 5C) and thus function as unusual flavoprotein dioxygenases that mediate ring dehydrogenation, CoA-ester oxygenolysis and a final stereoselective ring epoxidation^34^. To examine the roles of TpoE and TpoD in the biosynthesis of 3,7-dHT in *Pseudomonas* spp., we heterologously produced and purified these enzymes. Similarly, enzymes from the early PAA catabolism (PaaABCE, PaaG and PaaZ-E256Q) were obtained and employed for the biochemical generation of **3**^27,29,31^, which represents the likely substrate for TpoE. Following enzymatic formation of **3**^34^, the assays were complemented by TpoE, TpoD or a combination thereof. Interestingly, while the addition of TpoD had no effect, TpoE converted **3** into a new compound, whose mass and spectroscopic properties were consistent with the formation of epoxytropone-2-carboxyl-CoA **6**, resulting from ring dehydrogenation and subsequent epoxidation (Figure 6, Figure S10). Surprisingly, in contrast to previous observations with the homolog TdaE^Pi^ from *P. inhibens*^34^, TpoE appeared not to cleave the thioester itself. This implied the necessity for an additional thioesterase functionality en route to 3,7-dHT, for which TpoD is an evident prime candidate. Indeed, when both enzymes were added together, **7** was produced, as verified by LC-MS analysis and comparison to the identical TdaE-produced compound (Figures S10-S11). Taken together, these assays establish the roles of both TpoD and TpoE by showing that they jointly produce the highly reactive **7**, which subsequently degrades to tropolone upon spontaneous decarboxylation. Further studies will be required to identify the enzymes for the final steps in 3,7-dHT biosynthesis, as no apparent candidates for these ring hydroxylation steps are encoded in the *Pseudomonas* BGCs.

**Figure 6.**
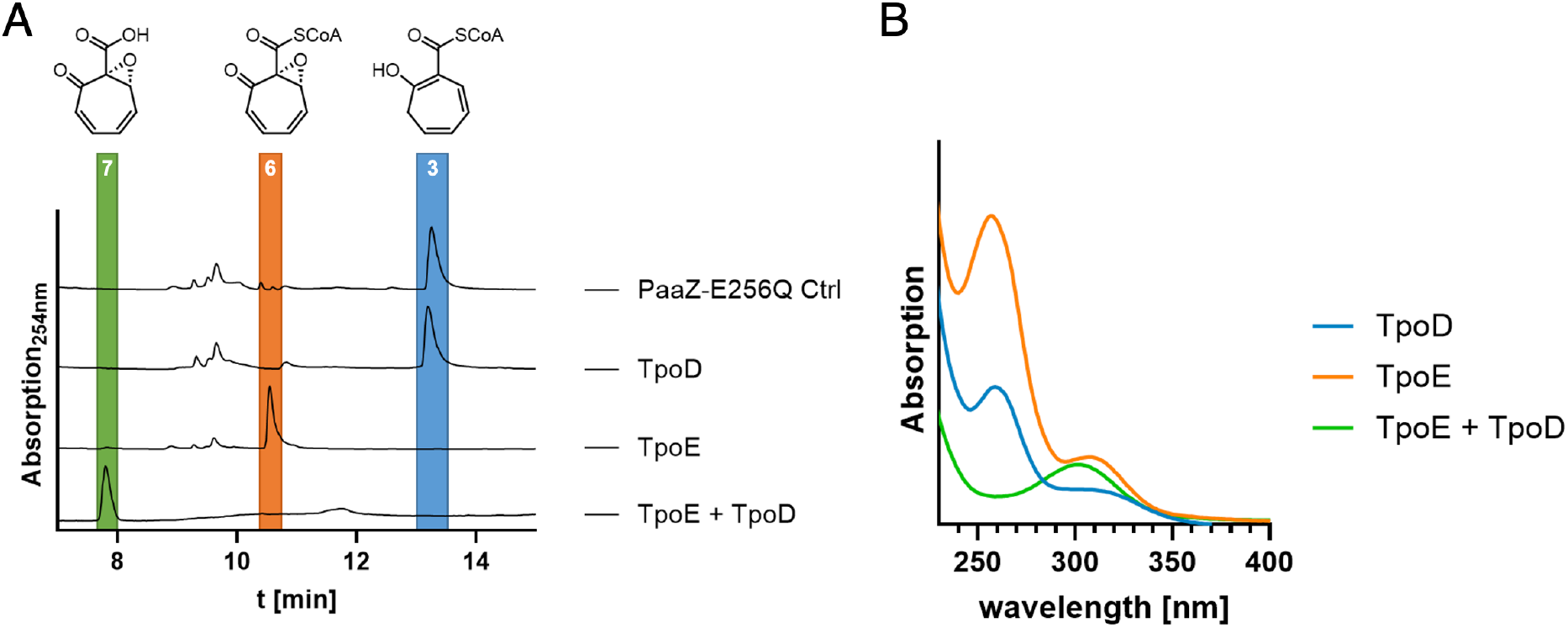
HPLC analysis of biochemical assays with TpoD and TpoE. A. The PaaZ-E256Q control (Crtl) variant with deficient aldehyde dehydrogenase domain was used to generate 3-oxo-5,6-dehydrosuberoyl-CoA semialdehyde, which undergoes Knoevenagel condensation to yield **3**, highlighted in blue. This precursor was incubated for 15 min with 1 µM of TpoD and/or TpoE. For the traces PaaZ-E256Q Ctrl, TpoD and TpoE the aqueous phase is shown (no significant amounts of any intermediate could be observed in the organic phase; not shown), for trace TpoE + TpoD the organic phase is shown (no significant amounts of any intermediate could be observed in the aqueous phase; not shown). B. UV-visible spectra of the highlighted compounds showing the absence of coenzyme A absorbance at 260 nm for compound **7**.

### Comparative analysis of tropolone BGCs

Given the essential role of the majority of the *tpo* biosynthetic genes, we assessed the presence and diversity of this non-canonical class of BGC across sequenced bacteria, especially as diverse tropolones are known to be produced across multiple genera. Due to the conserved requirement for TpoE/TdaE-like flavoproteins across diverse tropolone biosynthetic pathways^34^, we therefore used TpoE, TdaE^Pi^ and TdaE^Bp^ as bait proteins to identify further tropolone BGCs.

A strategy was devised to maximise the diversity of the resulting dataset. Here, the top 1000 BlastP hits for each TpoE/TdaE protein were obtained from the non-redundant NCBI protein database and filtered using a 95% identity cut-off to remove highly similar proteins and thus reduce the redundancy of associated BGCs. Putative BGCs were then retrieved as 35 kb regions centred on the *tpoE*/*tdaE* homologues. Following the removal of poor-quality sequences, the resulting set of 319 BGCs were grouped into 67 BGC families using BiG-SCAPE^58^ (Figures S12). Here, relatively permissive networking parameters were used due to the overall diversity of the dataset and the lack of multiple “anchor domains” that BiG-SCAPE usually uses for well-characterised classes of BGC. Diverse representatives of BGC families were then selected (86 clusters in total) for pan-family synteny analysis using clinker^59^ (Supplementary File 2, Figure S13). A subset of these BGCs with tropolone-like features are shown in Figure 7. An additional *Pseudomonas*-only analysis was performed to understand cluster distribution and synteny within this genus (Figure S14). Mapping these BGCs to *Pseudomonas* phylogeny (Figure S1) indicates that tropolone BGCs are rare but most commonly associated with the *Pseudomonas putida* group. Presence of the BGC in other *Pseudomonas* species groups supports some horizontal transfer within the genus.

**Figure 7.**
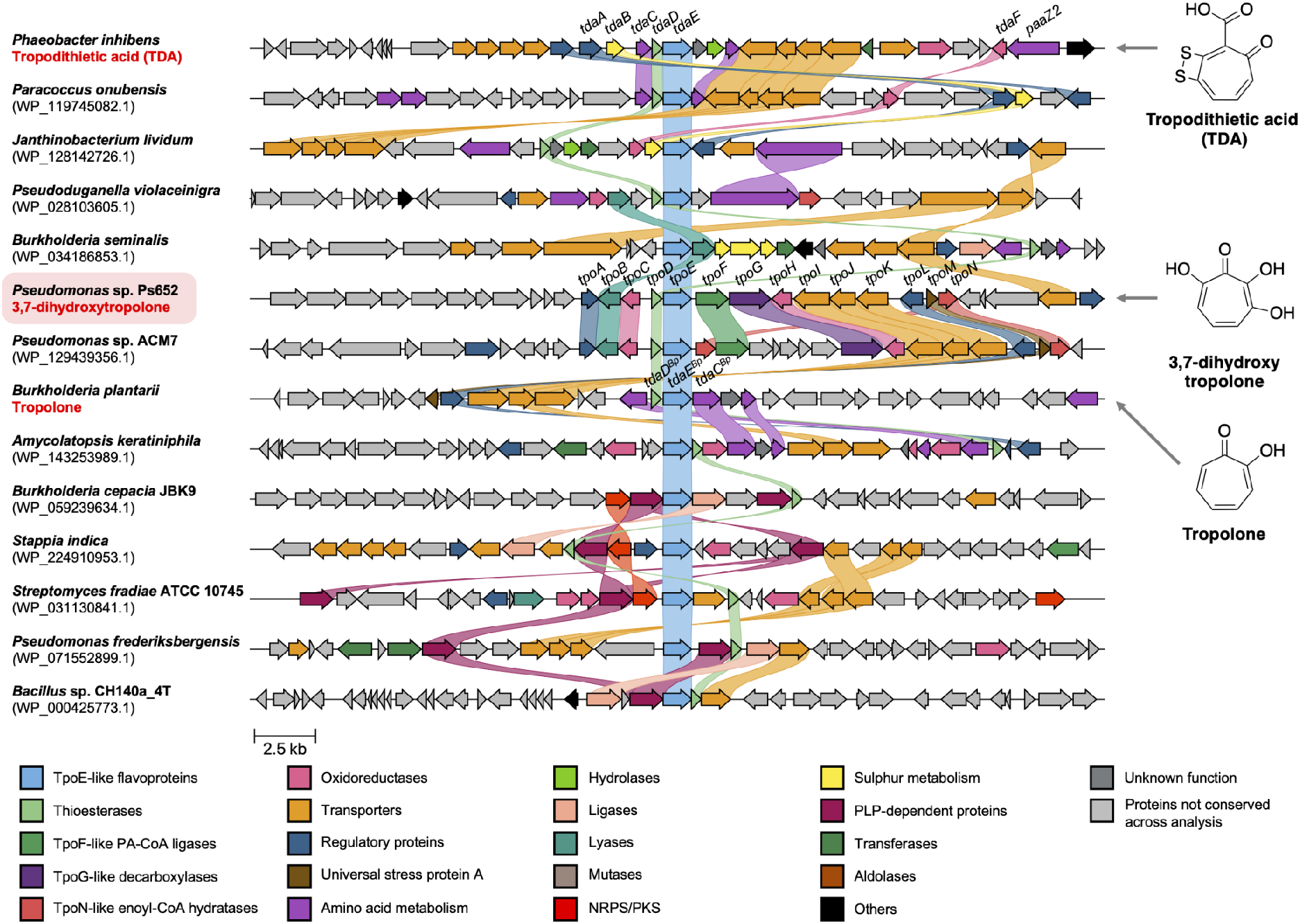
Comparison of bacterial tropolone BGCs and related BGCs identified by flavoprotein-led mining. BGC synteny is visualised using clinker^59^, where genes with at least 30% identity are colour-coded and linked. Previously characterised tropolone BGCs are highlighted, while the accessions of the TpoE homologues are listed for all other BGCs. Supplementary File 2 and Figures S13-S14 provide further BGC comparisons.

We were interested in assessing whether the presence of PAA catabolon genes is a common feature of these BGCs, given the importance of TpoF, TpoG and TpoN in 3,7-dHT biosynthesis (Figures 4A and 5C). *tpoF* and *tpoG* homologues are always clustered in *Pseudomonas* BGCs, where a subset of BGCs feature additional PAA catabolon genes next to *tpoF*, such as *Pseudomonas* sp. ACM7, which has an additional six PAA catabolon genes within the BGC (Figure S14). In contrast, TpoF and TpoG homologues are not encoded in BGCs from other genera, which instead encode a variety of alternative proteins that potentially boost precursor supply and activate the PAA catabolon, such as PaaZ2 in *P. inhibens* and *Janthinobacterium lividum*, or 3-deoxy-7-phosphoheptulonate synthase and prephenate dehydratase in *Burkholderia*, *Trinickia* and *Pseudoduganella*. This clustering of diverse PAA-associated genes with tropolone biosynthetic genes across varied BGCs suggests an uncommon form of convergent evolution that directs metabolic flux towards the biosynthesis of closely related specialised metabolites. Here, the tight regulation of the PAA catabolon could act as a selective pressure. Synteny across diverse BGCs also highlights potentially important roles for uncharacterised proteins, such as TpoB-like lyase/adolases (COG2321), which are conserved across BGCs in *Burkholderia*, *Serratia* and *Pseudoduganella*, or transporter complexes with homologues across multiple BGCs.

Thioesterase TpoD has an essential role in 3,7-dHT biosynthesis. In contrast, previous *in vitro* enzyme assays suggested that a functional thioesterase may not be strictly required in *Burkholderia* and *P. inhibens*^34^, even though the corresponding BGCs encode thioesterases (TdaD) with the same conserved domain (COG0824). Our genomic analysis demonstrated that TpoD/TdaD-like thioesterases are encoded in all BGCs that are closely related to characterised tropolone BGCs. In addition, a different family of small thioesterase-like proteins (COG5496) are also associated with a subset of uncharacterised clusters from diverse Gram-negative and Gram-positive bacteria (such as *Amycolatopsis keratiniphila*, *Burkholderia cepacia* and *Streptomyces fradiae*). These data hint at a conserved and important role for thioesterase-like proteins in tropolone biosynthesis, as well as providing lead BGCs for the discovery of novel troponoids^22,23^.

More broadly, the BiG-SCAPE and clinker datasets highlight the vast evolutionary diversity of BGCs associated with TpoE-like flavoproteins. It is likely that many BGCs in the broader dataset do not produce tropolones and are instead involved in alternative biosynthetic pathways, given the high frequency of diverse biosynthetic proteins encoded across clusters, including NRPSs, PKSs, transferases, amino acid metabolism proteins, ligases, aminotransferases and oxidoreductases (Figures 7 and S13). This clustering indicates that TpoE/TdaE homologues are reliable markers of uncharacterised BGCs involved in specialised metabolism.

### Conclusions

The tropolone chemophore is found in a variety of bioactive specialised metabolites^22,23^. These molecules have broad ecological relevance^60,61^ and medicinal promise, where some synthetic derivatives are under investigation as lead compounds for multiple diseases^62–65^. Here, we have demonstrated that the tropolone (*tpo*) BGC in *Pseudomonas* sp. Ps652 is responsible for the biosynthesis of both 7-HT and 3,7-dHT, which represents the first report of a 3,7-dHT BGC in a Gram-negative bacterium. 3,7-dHT is produced in similar amounts to 7-HT (Figure S4 and S8), which indicates that it could represent the true final product of the pathway, especially given that it is more active towards *S. scabies* (Figure 4A and S7). In addition to the iron-binding activity of both molecules^17^ (Figure S7), the broad biological activity of both 7-HT and 3,7-dHT^65^ indicates that both molecules could function to inhibit prokaryote and eukaryote organisms in competitive soil and plant microbial communities. Notably, the *tpo* BGC has been identified in multiple *Pseudomonas* strains via screens of large strain collections for organisms that inhibit plant pathogens^13,20,21,66–68^, highlighting the high potency of these molecules amongst complex microbial communities. Future efforts should focus on understanding the function of tropolones in natural plant and soil colonisation conditions, as well as the interplay with other BGCs and proteins encoded in tropolone producers^19,69^ (Figure 2). For example, recent work by Gram and colleagues has highlighted the impact of tropodithietic acid on *Phaeobacter* species and associated microbial communities^60,61^.

Based on mutational data and prior analyses of homologous BGCs in *Pseudomonas*, we propose a naming scheme for the *tpo* BGC (Figure 3A). The high similarity of the Ps652 BGC to other characterised *Pseudomonas* suggests that these strains have the capacity to also make 3,7-dHT, although this hypothesis needs to be tested. Using a mixture of metabolomics and reporter assays, we show that cluster-situated genes (*tpoFG*) generate PA-CoA, a key precursor for tropolone biosynthesis via the PAA catabolon, which is depressed by PA-CoA (Figure 5A). This interplay between the PAA catabolon and tropolone biosynthesis is well-established in other organisms^22,23^, but the tight clustering of PAA genes within a tropolone BGC is unusual in Gram-negative bacteria. Informatic analysis indicates that some *Pseudomonas* strains have additional PAA genes embedded within tropolone BGCs (Figure S12), which could further support metabolic flux towards tropolone biosynthesis. The involvement of the PAA catabolon in *Pseudomonas* biosynthesis of tropolones was highlighted in a recent study by Wang *et al*. conducted in parallel to our work^70^, which showed that multiple PAA catabolon genes are essential for 7-HT production in *Pseudomonas donghuensis* HYS. Further work is required to understand how the hydroxyl groups of 7-HT and 3,7-dHT are introduced. Characterised tropolone hydroxylases are flavoproteins^23^, but there are no equivalent proteins in the *tpo* BGC. The BGC does encode two uncharacterised oxidoreductases (Figure 3A) that may be somehow involved in these biosynthetic steps.

As in other tropolone pathways, our data are consistent with the proposal that PAA catabolism is intercepted via an enoyl-CoA dehydratase (TpoN), which generates a reactive intermediate that is converted into the 7-membered tropolone precursor (**3**, Figure 5C). The previous cryptic role of an essential and conserved thioesterase (TpoD) could be elucidated in this work and involves the specific CoA-ester cleavage of **6** to afford **7** (Figure 6), which spontaneously decomposes to tropolone. This is consistent with a *lux* reporter assay (Figure 5A) showing that expression of the PAA catabolon is unaffected in the Δ*tpoD* mutant, which suggests that it functions downstream of the catabolon. The co-occurrence of thioesterases with TpoE-like flavoproteins in diverse BGCs (Figure 7) is consistent with both enzymes acting in consecutive pathway steps, although biochemical assays showed that the dehydrogenase in Burkholderia (TdaE^Bp^) can catalyse CoA cleavage itself by an unusual O_2_-dependent mechanism^34^. It will be interesting to see if TpoD homologues are essential in all tropolone biosynthetic pathways or whether TpoE homologues may in some cases suffice to ensure efficient CoA-ester cleavage. Finally, we showed that TpoE/TdaE homologues can be used to identify uncharacterised BGCs via targeted genome mining, which are strong candidates for the production of novel metabolites, such as the ketoacyl synthase associated BGCs (Figures 7, S12 and S13).

## EXPERIMENTAL

### Chemicals and reagents

Unless otherwise specified, all chemicals were obtained from Sigma-Aldrich (Merck). All solvents for extractions and chromatographic applications were supplied by Fisher Scientific. Synthetic 7-HT was provided by Prof. Ryan Murelli (Brooklyn College, The City University of New York). All enzymes were obtained from New England Biolabs (NEB) unless other specified. DNA purification kits were obtained from Qiagen. Ultrapure H_2_O was obtained using a Milli-Q purification system from Merck. Oligonucleotides were obtained (desalted) from Sigma-Aldrich and redissolved in Milli-Q H_2_O. All oligonucleotides used in this work are described in Table S2.

### Genome sequencing

A combined Illumina-Nanopore genome for *Pseudomonas* sp. Ps652 was obtained from MicrobesNG (Birmingham, UK). The strain was supplied for gDNA preparation as per MicrobesNG instructions. The supplied sequence was received as 5972118 bp in 5 contigs with 44.4x coverage. The genome was reordered using the Align & Reorder contigs tool in Mauve^71^ using *Pseudomonas* sp. SNU WT1 as a reference^72^. The exported reordered file was submitted to the MeDuSa server^73^ for scaffolding. The output scaffold produced was 5972218 bp in 3 contigs (5966768 bp, 5323 bp, and 127 bp). Contig 2 (127 bp) and contig 3 (5325 bp) were both found by BLAST^74^ to be part of contig 1 (5966768 bp). These were thus removed and 5966768 bp was used as the single scaffold genome, which was annotated using Prokka 1.13.3^75^ and submitted to the European Nucleotide Archive (Accession: OZ024668). The final genome has a size of 5.96 Mbp with 62.2% GC and 5371 genes. This assembly was submitted to antiSMASH 7.0.0 in July 2023 with “relaxed” detection strictness. The tropolone BGC will be deposited at MIBiG^76^ with accession BGC0002842.

### Strains and media

All strains and plasmids used in this study are described in Tables S1 and S3. NEB 5-alpha Competent *E. coli* cells (High Efficiency, C2987H, New England Biolabs) were used for all transformations of *E. coli* and performed as per manufacturer instructions. All media used are defined in Table S4, where all formulae are for 1 L of deionised water unless specified otherwise. All media were sterilised by autoclaving and standardised to pH 7.2 unless otherwise stated. All media ingredients were sourced from Sigma-Aldrich (Merck) unless otherwise stated. Antibiotics were added where necessary at the following concentrations: tetracycline (15μg/mL for *E. coli*, 25 μg/mL for *Pseudomonas* sp. Ps652); gentamicin (25 µg/mL for *E. coli*, 50 µg/mL for *Pseudomonas*). *E. coli* was grown in liquid LB medium at 30 °C or 37 °C, or LB agar at 37 °C for 16-20 hours. Stocks of *E. coli* strains were stored at −70 °C in 25% glycerol. *Pseudomonas* strains were grown in Lennox medium at 30 °C for 16-20 hours unless otherwise specified. *Pseudomonas* strains were stored at −70 °C as 925 μL overnight culture + 75 μL DMSO, or 500 μL overnight culture + 500 μL 50% glycerol. *S. scabies* 87-22 were stored as spore stocks that were grown to sporulation on instant mash agar (IMA) plates, harvested in 20% glycerol and stored at −70 °C. *P. infestans* strain 88069 (The Sainsbury Laboratory, UK) was maintained on rye sucrose agar (RSA). Where relevant, all plates were imaged using a Canon Ixus 175 digital camera.

### Antimicrobial assays with *Streptomyces scabies* 87-22

10 µL of a *S. scabies* 87-22 spore stock was resuspended in 1 mL Milli-Q H_2_O, before 35 µL of this mixture was spread thoroughly onto IMA plates using a sterile cotton bud. Fractions / extracts were dried *in vacuo* using a GeneVac and redissolved in 50-100 µL ethyl acetate and applied to filter paper disks, which were allowed to dry completely before being added to the IMA plate. Plates were incubated at 28 °C for 2 days or until *S. scabies* had grown sufficiently for biological activity to be clearly observed.

### Assays with *P. infestans* strain 88069

A 5 mm diameter plug was taken from the actively growing outer edge of *P. infestans* mycelium using a heat and ethanol sterilised number 3 metal corer, and placed in the centre of a standard petri dish on RSA. 15 μL of overnight liquid culture of the *Pseudomonas* strain to be tested was then spotted in duplicate 2.5 cm either side of the central plug. Spotted liquid culture was then dried in a biosafety level 2 cabinet, before the plate was transferred to 20 °C and incubated for 10 days.

### Iron Binding Assays

Iron binding was tested using chrome azurol S (CAS) agar plates as originally defined by Schwyn and Neilands^50^. The binding of CAS / hexadecyltrimethylammonium (HDTMA) in the medium complexes with ferric iron to produce a blue colour. Siderophores are able to compete for the binding of iron, liberating it from CAS/HDTMA, producing a colour change from blue to orange. 5 µL of overnight culture or 5 µg of purified compounds dissolved in MeOH were added as spots to the plate and allowed to dry. If overnight cultures were used, plates were incubated for 1 day at 30 °C before being imaged. For pure compounds, the solvent was allowed to dry, plates were then stored at room temperature overnight to allow for diffusion of applied molecules before being imaged.

### In-frame deletions of *Pseudomonas* genes

In-frame deletion mutants of *Pseudomonas* strains were constructed by markerless 2-step allelic exchange^77^. Mutant alleles were designed by selecting approximately 500 bp sequences upstream and downstream of the region to be deleted, in-frame with the gene(s) to be deleted. These were either amplified by PCR (primers in Table S2) and cloned into the pTS1 suicide vector^8^ by Gibson assembly, or synthesised by Twist Biosciences Ltd., where the genes were provided in the pTS1 vector. ^8^Gibson assembly was performed using Gibson Assembly® Master Mix or NEBuilder® HiFi DNA Assembly mix according to manufacturer instructions. The pTS1 plasmid containing the shortened, mutant allele was used to transform *Pseudomonas* sp. Ps652 as follows. Ps652 was streaked from glycerol stocks onto Lennox agar and grown at 30 °C until visible colonies appeared. Single colonies were used to inoculate 10 mL Lennox broth per transformation. Liquid cultures were grown overnight before being centrifuged at 4000 x g for 8 minutes and resuspended in sterile 300 mM sucrose (2 mL). 2 x 1 mL was transferred to two 2 mL microcentrifuge tubes and centrifuged at 11,000 x g for 1 minute and the supernatant was removed by decanting. 1 mL 300 mM sucrose was added and resuspended by gentle vortex mixing. This was centrifuged at 11,000 x g for 1 minute and the supernatant removed as before. This wash was repeated twice more, and the cells from both tubes resuspended in a total of 100 µL 300 mM sucrose. 2 µL of DNA was added to each 100 µL aliquot and mixed carefully with a pipette tip.

Samples were then added to electroporation cuvettes (2 mm) and electroporation was performed using an Eppendorf Eporator at a setting of 2500 V. 900 µL of Lennox broth was added immediately after electroporation and the samples were mixed by gentle pipetting. Electroporated cells were incubated at 30 °C for 1 hour with shaking at 250 rpm. 100 µL of electroporated cells was then plated on Lennox agar with antibiotics as required and incubated at 28 °C until colonies were visible. Individual colonies were inoculated into 10 mL Lennox broth and grown for 16-18 hours at 28 °C with 250 rpm shaking. These cultures were then diluted to 10^−6^ and plated on Lennox agar supplemented with 10% sucrose, and incubated at 28 °C until visible colonies formed. These colonies were restreaked and then checked by colony PCR with GoTaq 2X Master Mix for the mutant allele, with the parent strain as the control. Positive clones were restreaked again to single colonies, tested again, and then grown 16-18 hours at 28 °C with 250 rpm shaking in Lennox broth supplemented with tetracycline. 75 µL DMSO was added to 925 µL of culture and this was stored at −70 °C.

### Preparation of extracts from *tpo* mutants

All strains were streaked from stocks to single colonies on Lennox agar plates with selection if necessary, and grown overnight in biological triplicate in 5 mL MKBG in 50 mL centrifuge tubes with sterilised foam bungs and no selection to prevent background activity from added antibiotics. 5 mL EtOAc was added, and tubes were agitated for 1 hour at 250 rpm. These were then centrifuged at 4,000 x g for 5 minutes to pellet any cell debris and ensure proper separation of aqueous and organic phases. 1.5 mL samples of the organic phase were taken for each, and dried fully by centrifugation under reduced pressure at 30 °C on a Genevac EZ-2 Plus system using the ‘Low BP’ setting, before being redissolved in 1.5 mL 100% MeOH and stored at −30 °C until use.

### Complementation of mutants

Ps652 Δ*hcn*Δ*tpoD* was complemented by PCR amplification of *tpoD* and the entire intergenic region preceding it (primers in Table S2) and cloning into the multicopy replicative vector pME6032^48^ by Gibson assembly using Gibson Assembly® Master Mix or NEBuilder® HiFi DNA Assembly mix according to manufacturer instructions. Assembled constructs were transformed into NEB 5-alpha *E. coli* according to manufacturer instructions and described above, and plated on LB agar supplemented with 15 µg/mL tetracycline. Plasmids were isolated from replicate lines and clones positive for pME6032::*tpoD* were identified by PCR with prepared plasmid as template (primers in Table S2). Ps652 Δ*hcn*Δ*tpoD* was transformed with pME6032 or pME6032::*tpoD* by electroporation as described above for in-frame deletion mutants and plated on Lennox agar supplemented with 25 µg/mL tetracycline. Single colonies were inoculated into 10 mL Lennox broth and plasmids isolated, followed by diagnostic digest with BamHI and HindIII revealing an insert of the expected size. Positive clones were maintained as described above.

Both *tpoE* and *tpoF* were complemented using the pUC18-mini-Tn7-Gm gentamicin-selective vector (Table S3). In each case, the 5’-UTR of *tpoD* was PCR amplified along with *tpoE* / *tpoF* and cloned into the BamHI site of pUC18-mini-Tn7-Gm by Gibson assembly, with the *tpoD* 5’-UTR directly upstream of *tpoE* / *tpoF* to allow native regulation. Assembled constructs were transformed into NEB 5-alpha *E. coli* cells as described above with selection on 25 µg/mL gentamicin. Plasmids were isolated from replicate lines and screened by PCR amplification of the whole insert (5’-UTR and *tpoE* / *tpoF*). Positive clones were maintained as described above. Ps652 Δ*hcn*Δ*tpoE* and Ps652 Δ*hcn*Δ*tpoF* were transformed with the corresponding constructs, along with the pTNS2 helper plasmid^51^, by electroporation as described above and selected on Lennox agar supplemented with 50 µg/mL gentamicin. Clones carrying the insertion were identified by PCR for the whole insert (primers in Table S2).

Chemical complementation was performed for Ps652 Δ*hcn*Δ*tpoE* and Ps652 Δ*hcn*Δ*tpoF* by preparing extracts as described above with 3 replicates per strain, with the addition of 1 mM phenylacetic acid (PAA) as appropriate to the MKBG medium.

### Liquid chromatography - mass spectrometry (LC-MS)

LC-MS/MS analysis was performed on a Q-Exactive mass spectrometry system (Thermo Fisher Scientific). A Luna Omega 1.6μm Polar C18 100Å (50 x 2.1mm) column (Phenomenex) was used with 0.1% formic acid in H_2_O and MeOH as the mobile phase. Gradients were linear 0-95% MeOH from 1 to 6 minutes, followed by 95% MeOH until 8.8 minutes to wash off any material still associated with the column. Spectra were acquired in either positive or negative mode using a scan range of *m/z* 50-500. Mass spectra were obtained in positive mode using full MS/dd-MS^2^ acquisition settings with the following specific parameters: chromatography peak width = 7s; Full MS settings: resolution = 70,000, AGC target = 3×10^6^, maximum IT = 100 ms, scan range 50 to 500 *m/z*; dd-MS^2^ settings: resolution 17,500, AGC target = 1×10^5^, maximum IT = 50 ms, loop count = 5, isolation window 1.5 *m/z*, (N)CE/ stepped nce 20, 40, 60; dd settings: minimum AGC target = 8×10^3^, exclude isotopes ON, dynamic exclusion = 1s. Additionally for some runs, single ion monitoring (SIM) was performed in parallel with the data acquisition previously described using the following parameters: resolution = 70,000, AGC target = 5×10^4^, maximum IT = 200 ms, scan range 150 to 2000 *m/z*, isolation window 1.5 *m/z,* inclusion masses *m/z* 139.10140 (M+H, positive) and *m/z*155.03360 (M+H, positive).

### Production and extraction of 3,7-dihydroxytropolone

To avoid reisolating pyoverdine when using UV absorption to track tropolones throughout purification, a pyoverdine-null mutant (Ps652 Δ*hcn*Δ*pyo*) was used for production and purification of tropolones. Ps652 Δ*hcn*Δ*pyo* was grown for 16 hours in 2 × 10 mL MKBG in plastic universal 30 mL tubes. This culture was then used to inoculate 2 L of MKBG in 200 × 10 mL aliquots in 50 mL centrifuge tubes with foam bungs, using 100 µL seed culture per 10 mL, and then grown for 24 hours at 30 °C with shaking at 250 rpm. Culture from all 200 tubes was then combined into a single 2 L volume. Cells were removed by centrifugation at 8,000 x g for 8 minutes at 4 °C in a Sorvall Lynx 6000 centrifuge (Thermo Fisher Scientific). The supernatant was collected and cells discarded. The supernatant was concentrated by rotary evaporation to a volume of 1 L. Following the method of Jiang *et al.*^17^, the supernatant was then extracted 3 times with 0.5 L of EtOAc before being acidified to pH 2 using HCl. The aqueous phase was then extracted 3 more times with 0.5 L to yield a total of 3 litres of organic extract. All organic extracts were combined and dried by rotary evaporation.

### Purification of 3,7-dihydroxytropolone

The dried organic extract was dissolved in 3 mL MeOH and subjected to size-exclusion chromatography using a Sephadex® LH-20 column on an ÄKTA Pure system (Cytiva) in 100% MeOH attached to an Optilab® differential refractive index detector (Wyatt Technology). Flow rate was 1 mL / min, and 10 mL fractions were collected from 350 min to 700 min. UV absorbance was monitored at 330 nm to detect tropolones. The 3 mL of organic extracted was processed in 2 injections. All fractions were dried on a Genevac EZ-2 Plus system using the ‘Low BP’ setting at 30 °C, and all fractions with strong absorbance at 330 nm were redissolved in MeOH, diluted 1:5, and 5 µL analysed by LC-MS on a Q-Exactive Orbitrap mass spectrometer (Thermo Fisher). Several fractions showed high levels of 7-hydroxytropolone, 3,7-dihydroxytropolone, or a mixture of both. These fractions were selected for further purification.

Fractions containing hydroxylated tropolones were redissolved in 100% MeOH and fractions with similar LC-MS profiles were combined. These fractions were processed by semi-preparative HPLC on a Dionex Ultimate 3000 system (Thermo Fisher Scientific) with a Luna Omega 5 µm Polar C18 100Å 250×10 mm column (Phenomenex) in 5 injections. A multistep gradient of H_2_O + 0.5% formic acid and MeOH + 0.5% formic acid was used (see Table S7 for details) with a flow rate of 4 mL/min and UV absorbance monitored at 254 nm and 327 nm. The chromatogram showed two clear peaks at 327 nm, with the earlier peak representing 3,7-dihydroxytropolone and the later peak representing the 7-hydroxytropolone (Figure S4). All fractions corresponding to 3,7-dihydroxytropolone were combined and dried overnight on a Genevac EZ-2 Plus system using the ‘HPLC Lyo’ setting at 30 °C. 2.8 mg of 3,7-dihydroxytropolone was obtained and was observed to be a white powder with a slight pink hue.

### NMR analysis of 3,7-dihydroxytropolone

3,7-dihydroxytropolone (2.8 mg) was redissolved in 500 µL CD_3_OD and shaken for 10 minutes to exchange OH protons for deuterium, and dried on a Genevac EZ-2 Plus system on the ‘Low BP’ setting at 30 °C. The sample was then again redissolved in 500 µL CD_3_OD and analysed on a Bruker AVANCE NEO 600 MHz Spectrometer equipped with a TCI cryoprobe. The experiments were carried out at 298 K with the residual CD_3_OD solvent used as an internal standard (δ_H_/δ_C_ 3.31/49.0 ppm). Resonances were assigned through 1D ^1^H and ^13^C experiments. Spectra were analysed using Bruker TopSpin 3.5 software. NMR spectra are reported in Figures S5-S6. ^1^H NMR ∂= 6.98 (3H, m). ^13^C NMR ∂= 119.24 (2C), 129.44, 158.11 (2C), 158.94 (2C).

### Reporter assay for PAA catabolon

The 5’-UTR of *paaA* was PCR amplified (primers in Table S2) and cloned by Gibson assembly into the BamHI to XcmI site of pUC18-mini-Tn7-Gm-lux plasmid upstream of *luxC*. Assembled constructs were transformed into NEB 5-alpha *E. coli* as described above and plated on LB agar supplemented with 25 µg/mL gentamicin. Plasmids were isolated from replicate lines, amplified by PCR (primers in Table S2) and verified by Sanger sequencing (Eurofins). In order to optimise the distance between the ribosome binding site and the start of *luxC*, site-directed mutagenesis via PCR (Table S2) was performed using Phire Hot Start II DNA Polymerase (Thermo Fisher Scientific) and again verified by Sanger sequencing. pUC18-mini-Tn7-Gm::paa-lux was transformed, along with the pTNS2 helper plasmid, into the relevant Ps652 strains by electroporation as described above and selected on gentamicin. Positive clones were identified by PCR screening (primers in Table S2).

For luminescence measurements, all strains were inoculated from single colonies into 10 mL MKBG medium with or without addition of 1 mM PAA and grown for 12 hours at 30 °C with shaking at 250 rpm. All cultures were then diluted to an optical density at 600 nm of 1.0 in the relevant medium in 1.5 mL microcentrifuge tubes per strain (5 replicates) and shaken for 1 hour at 30 °C. Luminescence measurements were then taken on a GloMax®-Multi Jr instrument (Promega).

### Analysis of flavoprotein-associated BGCs

The flavoproteins from *Pseudomonas* sp. Ps652 (TpoE, VVM49077.1), *Burkholderia plantarii* (TdaE^Bp^, WP_042624079.1) and *Phaeobacter inhibens* (TdaE^Pi^, WP_014881725.1) were used as queries for BlastP searches using the NCBI non-redundant protein database with default search paraments^74^. The top 1,000 hits were retrieved from each search (3,000 accessions in total), which were then filtered for duplicates to 1,543 accessions. This list of proteins was then filtered using a 95% sequence identity cut-off using CD-Hit^78^ to reduce redundancy in downstream analyses to provide a list of 526 protein accessions whose identity to each other was lower than 95%. This accession list was then used to retrieve 35 kb Genbank files centred on the input accessions. The dataset was further filtered to remove poor quality output files that resulted from poor sequencing data, such as from metagenome-assembled genomes and contig edges of isolated genomes. Additional output files from non-bacterial sources were also removed. The final output consisted of 319 putative BGCs as Genbank files (Supplementary File 1).

The Genbank files were each edited to make them compatible with features within BiG-SCAPE^58^ 1.1.5 (glocal mode and anchor domains) by adding a `/product=“other”` qualifier following the region feature, and a `/gene_kind=“biosynthetic”` qualifier to the CDS entry for the TdaE/TdoE homologue in each file. BiG-SCAPE 1.1.5 was run using the following parameters with the 319 BGCs plus the tropodithietic acid BGC (MiBIG BGC0000932):

`--mix --mode glocal --cutoffs 0.5 0.75 0.85 --anchorfile`
`anchor_domains.txt --include_singletons --clans-off`

The anchor file included the following information to anchor the BGC analysis on the central flavoprotein of each cluster:

`PF00441 Dehydrogenase_C-term [Others]`
`PF02771 Dehydrogenase_N-term [Others]`

Manual assessment determined that a cut-off of 0.75 separated clusters into meaningful families without overly fragmenting the output into a large proportion of singletons. In total, the clusters were grouped into 67 families, which were further visualised and annotated using Cytoscape 3.10.1 (Figure S12).

To visualise the synteny and diversity of the resulting families, one or two representative clusters were selected from each family and visualised using clinker^59^ 0.0.27 (Figure 7, Figure S13 and Supplementary File 2).

### Analysis of *Pseudomonas* phylogeny and tropolone BGC synteny

TpoE (VVM49077.1) was used as a query for BlastP searches using only *Pseudomonas* sequences within the NCBI non-redundant protein database and with a 50% identity cut-off. 22 proteins were retrieved, which were then filtered using a 99% sequence identity cut-off using CD-Hit^78^ to provide 8 protein accessions. 35 kb Genbank files were obtained that were centred on each protein accession. BGC synteny was then visualised using clinker^59^ 0.0.27 (Figure S12). To assess the taxonomy of Ps652 and the associated distribution of tropolone BGCs, the genomes of Ps652 and the 7 additional strains were used as input for AutoMLST^35^. Default AutoMLST settings were used plus ModelFinder and IQ-Tree bootstrapping. AutoMLST automatically picks reference strains (up to a maximum of 50 strains for the tree) for remainder of the tree and associated conserved genes for multi-locus phylogenetic analysis. The precise strains used for the tree were manually adjusted to reduce taxonomic redundancy and obtain broader representation of the *Pseudomonas* genus. The conserved genes selected by AutoMLST for tree building are listed in Supplementary File 1. In addition to the 8 input strains, the other strains in the tree were assessed for the presence of a tropolone BGC. The resulting tree was visualised using iTOL^79^ (Figure S1).

### Cloning and recombinant protein production of TpoE and TpoD

The genes *tpoE* and *tpoD* were synthesized and codon optimized for *E. coli* by Biocat (Heidelberg, Germany). Using restriction enzyme digestion with NotI (3’) and NcoI (5’) both fragments were cloned in a modified pET28b-vector, which adds an N-terminal His6x-gb1-Tag to the recombinant protein. This vector contained a hexahistidine-tag for immobilized metal-affinity chromatography (IMAC) purification, a solubility enhancer gb1-tag (B1 domain of *Streptococcal* protein g) and a TEV protease cleavage-site. Correct cloning was confirmed by Sanger sequencing.

The resulting constructs were transformed in *E. coli* BL21 (DE3) pL1SL2 cells ^80^. These cells contain a GroES/GroEL chaperonin system, which helps with protein folding. Single colonies were picked, and starter cultures were grown in LB-medium supplemented with 50 µg/mL kanamycin, 100 µg/mL ampicillin and 20 µg/mL chloramphenicol at 37 °C and 130 rpm overnight. Main cultures with TB medium and the same amount of antibiotics were inoculated with 1% of the overnight cultures and grown at 37°C and 130 rpm to an OD_600nm_ between 0.4 – 0.6. After that, protein expression was induced with 500 µM IPTG. The cultures were subsequently incubated at 18 °C and 130 rpm until the following day. The cultures were harvested by centrifugation at 5,000 x g for 30 min at 4 °C, resuspended in 0.9% NaCl (to remove residual medium), centrifuged again with the same parameters and subsequently frozen as pellet at −20 °C.

### Protein purification of TpoE and TpoD

The frozen cell pellets were resuspended in lysis buffer containing 300 mM NaCl, 10% glycerol (v/v), 50 mM sodium phosphate buffer (Na_2_HPO_4_/NaH_2_PO_4_) at pH 7.4. The resuspended cells were lysed by ultrasonication (3 s pulse, 3 s pause, 4 min pulse time, amplitude: 60%, repeated once). The lysate was cleared by centrifugation (18,000 x g, 30 min and 4 °C) and sterile filtration (pore size 0.22 µm) and applied to a Ni-NTA column (Cytiva). The column was washed with lysis buffer containing an additional 30 mM imidazole to remove other protein impurities. Elution of the target protein was then carried out with lysis buffer supplemented with 500 µM imidazole. Subsequently, a desalting column was used to exchange buffer and remove imidazole. The obtained fractions were concentrated with an Amicon concentrator (Merck Millipore) and flash frozen in liquid nitrogen upon storage at −80 °C.

### Recombinant protein production and purification of PaaABCE, PaaG and PaaZ-E256Q

Enzymes from the PAA catabolon for the *in vitro* formation of tropolone precursor **3** were produced and purified as previously described ^27,29,31^.

### Enzymatic synthesis of 7

Compound **7**, the precursor for tropolone formation, was synthesized using the same substrate, enzymes and reaction conditions as previously described^34^.

### Turnover assay with TpoE and TpoD

To the substrate mix containing approximately 0.5 mM **4** (enzymatically produced by PaaABCE, PaaG & PaaZ-E256Q) in Tris-HCl 50 mM buffer either 1 µM of TpoE, 1 µM of TpoD or 1 µM of both enzymes was added. The samples were put on a tabletop shaker in Eppendorf tubes with open lids and incubated for 15 min at 30 °C and vigorous shaking (900 rpm). Afterwards, the reaction was quenched by adding an equal volume of ethyl acetate with 0.1% formic acid and thorough vortexing of the samples. The samples were centrifuged at 18,000 x g for 10 min and the organic phase was separated from the aqueous phase. The organic ethylacetate phase was dried under N_2_ gas flow and the samples were resuspended in acetonitrile.

### RP-HPLC-MS analysis of the enzyme assays

Organic and aqueous phases obtained from TpoE/TpoD assays were both analyzed by RP-HPLC on a Shimadzu LCMS-8030 Triple Quad Mass Spectrometer. A Sunfire C18 column (150 x 3 mm ID, 3.5 µM, Waters) with a guard column was used. For aqueous samples, 10 mM ammonium acetate buffer pH 4.5 was used as solution A1, for organic samples water + 0.1% formic acid (A2) was used. Solution B1 was acetonitrile and B2 acetonitrile + 0.1% formic acid. The HPLC program for both aqueous and organic phases was the same. The flow rate was set to 0.4 mL/min and the following gradient was used:

2 - 12% B (0 – 4 min), 12 % B (4 – 6 min), 12 – 60% B (6 – 14 min), 60% B (14 – 18 min), 60 – 100% B (18 – 19 min), 100% B (19 – 22 min), 100 – 2% B (22 – 23 min), 2% B (23 – 26 min).

Absorption was measured from 190 – 800 nm. For MS measurements, electrospray ionization (ESI) was used in both negative and positive mode with the following settings: 3 kV capillary voltage, 400 °C heat block temperature, 250 °C DL temperature and 3 L/min nebulizing gas flow.

## ACKNOWLEDGEMENTS

This work was funded by a UK Research and Innovation (UKRI) Biotechnology and Biological Sciences Research Council (BBSRC) Norwich Research Park Doctoral Training Partnership grant (BB/M011216/1) for A.D.M. The work was also supported by BBSRC Institute Strategic Programme grants (BBS/E/J/000PR9797, BBS/E/J/000PR9790, BBS/E/JI/230001C and BB/X01097X/1) for the John Innes Centre (JIC). We are very grateful for the technical assistance at JIC provided by Dr Lionel Hill for LC-MS, Dr Martin Rejzek for HPLC, Dr Govind Chandra for informatics and Dr Sergey Nepogodiev for NMR. We are thankful to Dr Natalia Miguel-Vior (JIC) for assistance with LC-MS data and helpful discussions, and to Prof. Ryan Murelli (Brooklyn College, The City University of New York) for sharing synthetic 7-HT. This work was further supported by the Swiss National Science Foundation (SNSF) by grant 212747 awarded to R.T.

## REFERENCES

1. Haas, D. & Défago, G. Biological control of soil-borne pathogens by fluorescent pseudomonads. Nat. Rev. Micro. 3, 307–319 (2005).

2. Yi, B. & Dalpke, A. H. Revisiting the intrageneric structure of the genus Pseudomonas with complete whole genome sequence information: Insights into diversity and pathogen-related genetic determinants. Infect. Genet. Evol. 97, 105183 (2022).

3. Hesse, C., Schulz, F., Bull, C. T., Shaffer, B. T., Yan, Q., Shapiro, N., Hassan, K. A., Varghese, N., Elbourne, L. D. H., Paulsen, I. T., Kyrpides, N., Woyke, T. & Loper, J. E. Genome-based evolutionary history of Pseudomonas spp. Environ. Microbiol. 20, 2142–2159 (2018).

4. Gross, H. & Loper, J. E. Genomics of secondary metabolite production by Pseudomonas spp. Nat. Prod. Rep. 26, 1408–1446 (2009).

5. Masschelein, J., Jenner, M. & Challis, G. L. Antibiotics from Gram-negative bacteria: a comprehensive overview and selected biosynthetic highlights. Nat. Prod. Rep. 34, 712–783 (2017).

6. Santos-Aberturas, J. & Vior, N. M. Beyond Soil-Dwelling Actinobacteria: Fantastic Antibiotics and Where to Find Them. Antibiotics 11, 195 (2022).

7. Cornelis, P. Iron uptake and metabolism in pseudomonads. Appl. Microbiol. Biotechnol. 86, 1637–1645 (2010).

8. Scott, T. A., Heine, D., Qin, Z. & Wilkinson, B. An L-threonine transaldolase is required for L-threo-β-hydroxy-α-amino acid assembly during obafluorin biosynthesis. Nat. Commun. 8, 15935 (2017).

9. El-Sayed, A. K., Hothersall, J., Cooper, S. M., Stephens, E., Simpson, T. J. & Thomas, C. M. Characterization of the Mupirocin Biosynthesis Gene Cluster from Pseudomonas fluorescens NCIMB 10586. Chem. Biol. 10, 419–430 (2003).

10. Biessy, A., Novinscak, A., Blom, J., Léger, G., Thomashow, L. S., Cazorla, F. M., Josic, D. & Filion, M. Diversity of phytobeneficial traits revealed by whole-genome analysis of worldwide-isolated phenazine-producing Pseudomonas spp. Environ. Microbiol. 21, 437–455 (2019).

11. Ossowicki, A., Jafra, S. & Garbeva, P. The antimicrobial volatile power of the rhizospheric isolate Pseudomonas donghuensis P482. PLoS ONE 12, e0174362 (2017).

12. Stringlis, I. A., Zhang, H., Pieterse, C. M. J., Bolton, M. D. & Jonge, R. de. Microbial small molecules - weapons of plant subversion. Nat. Prod. Rep. 35, 410–433 (2018).

13. Pacheco-Moreno, A., Stefanato, F. L., Ford, J. J., Trippel, C., Uszkoreit, S., Ferrafiat, L., Grenga, L., Dickens, R., Kelly, N., Kingdon, A. D., Ambrosetti, L., Nepogodiev, S. A., Findlay, K. C., Cheema, J., Trick, M., Chandra, G., Tomalin, G., Malone, J. G. & Truman, A. W. Pan-genome analysis identifies intersecting roles for Pseudomonas specialized metabolites in potato pathogen inhibition. eLife 10, e71900 (2021).

14. Köhl, J., Kolnaar, R. & Ravensberg, W. J. Mode of Action of Microbial Biological Control Agents Against Plant Diseases: Relevance Beyond Efficacy. Front. Plant Sci. 10, (2019).

15. Moffat, A. D., Elliston, A., Patron, N. J., Truman, A. W. & Lopez, J. A. C. A biofoundry workflow for the identification of genetic determinants of microbial growth inhibition. Synth. Biol. 6, ysab004 (2021).

16. Yu, X., Chen, M., Jiang, Z., Hu, Y. & Xie, Z. The two-component regulators GacS and GacA positively regulate a nonfluorescent siderophore through the Gac/Rsm signaling cascade in high-siderophore-yielding Pseudomonas sp. strain HYS. J. Bacteriol. 196, 3259–3270 (2014).

17. Jiang, Z., Chen, M., Yu, X. & Xie, Z. 7-Hydroxytropolone produced and utilized as an iron-scavenger by Pseudomonas donghuensis. Biometals 29, 817–826 (2016).

18. Chen, M., Wang, P. & Xie, Z. A Complex Mechanism Involving LysR and TetR/AcrR That Regulates Iron Scavenger Biosynthesis in Pseudomonas donghuensis HYS. J. Bacteriol. 200, 592 (2018).

19. Krzyżanowska, D. M., Ossowicki, A., Rajewska, M., Maciąg, T., Jabłońska, M., Obuchowski, M., Heeb, S. & Jafra, S. When Genome-Based Approach Meets the “Old but Good”: Revealing Genes Involved in the Antibacterial Activity of Pseudomonas sp. P482 against Soft Rot Pathogens. Front. Microbiol. 7, 1415 (2016).

20. Muzio, F. M., Agaras, B. C., Masi, M., Tuzi, A., Evidente, A. & Valverde, C. 7-hydroxytropolone is the main metabolite responsible for the fungal antagonism of Pseudomonas donghuensis strain SVBP6. Environ. Microbiol. 22, 2550–2563 (2020).

21. Tao, X., Zhang, H., Gao, M., Li, M., Zhao, T. & Guan, X. Pseudomonas species isolated via high-throughput screening significantly protect cotton plants against verticillium wilt. AMB Express 10, 193 (2020).

22. Guo, H., Roman, D. & Beemelmanns, C. Tropolone natural products. Nat. Prod. Rep. 36, 1137–1155 (2019).

23. Duan, Y., Petzold, M., Saleem-Batcha, R. & Teufel, R. Bacterial Tropone Natural Products and Derivatives: Overview of their Biosynthesis, Bioactivities, Ecological Role and Biotechnological Potential. ChemBioChem 21, 2384–2407 (2020).

24. Lindberg, G. D., Larkin, J. M. & Whaley, H. A. Production of Tropolone by a Pseudomonas. J. Nat. Prod. 43, 592–594 (1980).

25. Cai, X., Shi, Y.-M., Pöhlmann, N., Revermann, O., Bahner, I., Pidot, S. J., Wesche, F., Lackner, H., Büchel, C., Kaiser, M., Richter, C., Schwalbe, H., Stinear, T. P., Zeeck, A. & Bode, H. B. Structure and Biosynthesis of Isatropolones, Bioactive Amine-Scavenging Fluorescent Natural Products from Streptomyces Gö66. Angew. Chem. Int. Ed. 56, 4945–4949 (2017).

26. Geng, H., Bruhn, J. B., Nielsen, K. F., Gram, L. & Belas, R. Genetic dissection of tropodithietic acid biosynthesis by marine roseobacters. Appl. Environ. Microbiol. 74, 1535–1545 (2008).

27. Teufel, R., Mascaraque, V., Ismail, W., Voss, M., Perera, J., Eisenreich, W., Haehnel, W. & Fuchs, G. Bacterial phenylalanine and phenylacetate catabolic pathway revealed. Proc. Natl. Acad. Sci. U.S.A. 107, 14390–14395 (2010).

28. Olivera, E. R., Miñambres, B., García, B., Muñiz, C., Moreno, M. A., Ferrández, A., Díaz, E., García, J. L. & Luengo, J. M. Molecular characterization of the phenylacetic acid catabolic pathway in Pseudomonas putida U: the phenylacetyl-CoA catabolon. Proc. Natl. Acad. Sci. U.S.A. 95, 6419–6424 (1998).

29. Teufel, R., Gantert, C., Voss, M., Eisenreich, W., Haehnel, W. & Fuchs, G. Studies on the mechanism of ring hydrolysis in phenylacetate degradation: a metabolic branching point. J. Biol. Chem. 286, 11021–11034 (2011).

30. Berger, M., Brock, N. L., Liesegang, H., Dogs, M., Preuth, I., Simon, M., Dickschat, J. S. & Brinkhoff, T. Genetic analysis of the upper phenylacetate catabolic pathway in the production of tropodithietic acid by Phaeobacter gallaeciensis. Appl. Environ. Microbiol. 78, 3539–3551 (2012).

31. Teufel, R., Friedrich, T. & Fuchs, G. An oxygenase that forms and deoxygenates toxic epoxide. Nature 483, 359–362 (2012).

32. Brock, N. L., Nikolay, A. & Dickschat, J. S. Biosynthesis of the antibiotic tropodithietic acid by the marine bacterium Phaeobacter inhibens. Chem. Commun. 50, 5487–5489 (2014).

33. Cane, D. E., Wu, Z. & Epp, J. E. V. Thiotropocin biosynthesis. Shikimate origin of a sulfur-containing tropolone derivative. J. Am. Chem. Soc. 114, 8479–8483 (2002).

34. Duan, Y., Toplak, M., Hou, A., Brock, N. L., Dickschat, J. S. & Teufel, R. A Flavoprotein Dioxygenase Steers Bacterial Tropone Biosynthesis via Coenzyme A-Ester Oxygenolysis and Ring Epoxidation. J. Am. Chem. Soc. 143, 10413–10421 (2021).

35. Alanjary, M., Steinke, K. & Ziemert, N. AutoMLST: an automated web server for generating multi-locus species trees highlighting natural product potential. Nucleic Acids Res. 47, W276–W282 (2019).

36. Richter, M. & Rosselló-Móra, R. Shifting the genomic gold standard for the prokaryotic species definition. Proc. Natl. Acad. Sci. U.S.A. 106, 19126–19131 (2009).

37. Peix, A., Ramírez-Bahena, M.-H. & Velázquez, E. The current status on the taxonomy of Pseudomonas revisited: An update. Infect. Genet. Evol. 57, 106–116 (2018).

38. Meier-Kolthoff, J. P., Carbasse, J. S., Peinado-Olarte, R. L. & Göker, M. TYGS and LPSN: a database tandem for fast and reliable genome-based classification and nomenclature of prokaryotes. Nucleic Acids Res. 50, D801–D807 (2021).

39. Girard, L., Lood, C., Höfte, M., Vandamme, P., Rokni-Zadeh, H., Noort, V. van, Lavigne, R. & Mot, R. D. The Ever-Expanding Pseudomonas Genus: Description of 43 New Species and Partition of the Pseudomonas putida Group. Microorganisms 9, 1766 (2021).

40. Blin, K., Shaw, S., Augustijn, H. E., Reitz, Z. L., Biermann, F., Alanjary, M., Fetter, A., Terlouw, B. R., Metcalf, W. W., Helfrich, E. J. N., van Wezel, G. P., Medema, M. H. & Weber, T. antiSMASH 7.0: new and improved predictions for detection, regulation, chemical structures and visualisation. Nucleic Acids Res. 51, W46–W50 (2023).

41. Johnston, I., Osborn, L. J., Markley, R. L., McManus, E. A., Kadam, A., Schultz, K. B., Nagajothi, N., Ahern, P. P., Brown, J. M. & Claesen, J. Identification of essential genes for Escherichia coli aryl polyene biosynthesis and function in biofilm formation. NPJ Biofilms Microbiomes 7, 56 (2021).

42. Choi, O., Kim, J., Kim, J.-G., Jeong, Y., Moon, J. S., Park, C. S. & Hwang, I. Pyrroloquinoline quinone is a plant growth promotion factor produced by Pseudomonas fluorescens B16. Plant Physiol. 146, 657–668 (2008).

43. Kretsch, A. M., Morgan, G. L., Acken, K. A., Barr, S. A. & Li, B. Pseudomonas Virulence Factor Pathway Synthesizes Autoinducers That Regulate the Secretome of a Pathogen. ACS Chem. Biol. 16, 501–509 (2021).

44. Carroll, L. M., Larralde, M., Fleck, J. S., Ponnudurai, R., Milanese, A., Cappio, E. & Zeller, G. Accurate de novo identification of biosynthetic gene clusters with GECCO. BioRxiv 2021.05.03.442509 (2021). doi:10.1101/2021.05.03.442509

45. Ramette, A., Frapolli, M., Défago, G. & Moënne-Loccoz, Y. Phylogeny of HCN Synthase-Encoding hcnBC Genes in Biocontrol Fluorescent Pseudomonads and Its Relationship with Host Plant Species and HCN Synthesis Ability. Mol. Plant-Microbe Interact. 16, 525–535 (2003).

46. Carrión, O., Curson, A. R. J., Kumaresan, D., Fu, Y., Lang, A. S., Mercadé, E. & Todd, J. D. A novel pathway producing dimethylsulphide in bacteria is widespread in soil environments. Nat. Commun. 6, 6579 (2015).

47. McClerklin, S. A., Lee, S. G., Harper, C. P., Nwumeh, R., Jez, J. M. & Kunkel, B. N. Indole-3-acetaldehyde dehydrogenase-dependent auxin synthesis contributes to virulence of Pseudomonas syringae strain DC3000. PLoS Pathog. 14, e1006811 (2018).

48. Heeb, S., Itoh, Y., Nishijyo, T., Schnider, U., Keel, C., Wade, J., Walsh, U., O’Gara, F. & Haas, D. Small, stable shuttle vectors based on the minimal pVS1 replicon for use in gram-negative, plant-associated bacteria. Mol. Plant-Microbe Interact. 13, 232–237 (2000).

49. Takeshita, H., Mori, A., Kusaba, T. & Watanabe, H. Preparation of Polyacetoxytropones and Polyhydroxytropolones by Acetolysis and Hydrolysis of Halotroponoids by Acetyl Trifluoroacetate with Exhaustive Displacement of Halogens on the Tropone Ring. Predominant Formation of Reductive Acetolysates from Fully-Substituted Tropones. Bull. Chem. Soc. Jpn. 60, 4325–4333 (1987).

50. Schwyn, B. & Neilands, J. B. Universal chemical assay for the detection and determination of siderophores. Anal. Biochem. 160, 47–56 (1987).

51. Choi, K.-H., Gaynor, J. B., White, K. G., Lopez, C., Bosio, C. M., Karkhoff-Schweizer, R. R. & Schweizer, H. P. A Tn7-based broad-range bacterial cloning and expression system. Nat. Meth. 2, 443–448 (2005).

52. Fernández, C., Díaz, E. & Garcia, J. L. Insights on the regulation of the phenylacetate degradation pathway from Escherichia coli. Environ. Microbiol. Rep. 6, 239–250 (2014).

53. Chen, X., Xu, M., Lü, J., Xu, J., Wang, Y., Lin, S., Deng, Z., Tao, M. & Vieille, C. Biosynthesis of Tropolones in Streptomyces spp.: Interweaving Biosynthesis and Degradation of Phenylacetic Acid and Hydroxylations on the Tropone Ring. Appl. Environ. Microbiol. 84, e00349–18 (2018).

54. Spaepen, S., Versées, W., Gocke, D., Pohl, M., Steyaert, J. & Vanderleyden, J. Characterization of Phenylpyruvate Decarboxylase, Involved in Auxin Production of Azospirillum brasilense. J. Bacteriol. 189, 7626–7633 (2007).

55. Koma, D., Yamanaka, H., Moriyoshi, K., Ohmoto, T. & Sakai, K. Production of aromatic compounds by metabolically engineered Escherichia coli with an expanded shikimate pathway. Appl. Environ. Microbiol. 78, 6203–6216 (2012).

56. Crabo, A. G., Singh, B., Nguyen, T., Emami, S., Gassner, G. T. & Sazinsky, M. H. Structure and biochemistry of phenylacetaldehyde dehydrogenase from the Pseudomonas putida S12 styrene catabolic pathway. Arch. Biochem. Biophys. 616, 47–58 (2017).

57. Glassing, A. & Lewis, T. A. An improved Tn7-lux reporter for broad host range, chromosomally-integrated promoter fusions in Gram-negative bacteria. J. Microbiol. Methods 118, 75–77 (2015).

58. Navarro-Muñoz, J. C., Selem-Mojica, N., Mullowney, M. W., Kautsar, S. A., Tryon, J. H., Parkinson, E. I., Santos, E. L. C. de L., Yeong, M., Cruz-Morales, P., Abubucker, S., Roeters, A., Lokhorst, W., Fernandez-Guerra, A., Cappelini, L. T. D., Goering, A. W., Thomson, R. J., Metcalf, W. W., Kelleher, N. L., Barona-Gómez, F. & Medema, M. H. A computational framework to explore large-scale biosynthetic diversity. Nat. Chem. Biol. 16, 60–68 (2020).

59. Gilchrist, C. L. M. & Chooi, Y.-H. clinker & clustermap.js: automatic generation of gene cluster comparison figures. Bioinformatics 37, 2473–2475 (2021).

60. Lindqvist, L. L., Jarmusch, S. A., Sonnenschein, E. C., Strube, M. L., Kim, J., Nielsen, M. W., Kempen, P. J., Schoof, E. M., Zhang, S.-D. & Gram, L. Tropodithietic Acid, a Multifunctional Antimicrobial, Facilitates Adaption and Colonization of the Producer, Phaeobacter piscinae. mSphere 8, e00517–22 (2023).

61. Henriksen, N. N. S. E., Schostag, M. D., Balder, S. R., Bech, P. K., Strube, M. L., Sonnenschein, E. C. & Gram, L. The ability of Phaeobacter inhibens to produce tropodithietic acid influences the community dynamics of a microalgal microbiome. ISME Commun. 2, 109 (2022).

62. Hirsch, D. R., Schiavone, D. V., Berkowitz, A. J., Morrison, L. A., Masaoka, T., Wilson, J. A., Lomonosova, E., Zhao, H., Patel, B. S., Datla, S. H., Hoft, S. G., Majidi, S. J., Pal, R. K., Gallicchio, E., Tang, L., Tavis, J. E., Grice, S. F. J. L., Beutler, J. A. & Murelli, R. P. Synthesis and biological assessment of 3,7-dihydroxytropolones. Org. Biomol. Chem. 16, 62–69 (2018).

63. Cao, F., Orth, C., Donlin, M. J., Adegboyega, P., Meyers, M. J., Murelli, R. P., Elagawany, M., Elgendy, B. & Tavis, J. E. Synthesis and Evaluation of Troponoids as a New Class of Antibiotics. ACS Omega 3, 15125–15133 (2018).

64. Ireland, P. J., Tavis, J. E., D’Erasmo, M. P., Hirsch, D. R., Murelli, R. P., Cadiz, M. M., Patel, B. S., Gupta, A. K., Edwards, T. C., Korom, M., Moran, E. A. & Morrison, L. A. Synthetic α-Hydroxytropolones Inhibit Replication of Wild-Type and Acyclovir-Resistant Herpes Simplex Viruses. Antimicrob. Agents Chemother. 60, 2140–2149 (2016).

65. Meck, C., D’Erasmo, M. P., Hirsch, D. R. & Murelli, R. P. The biology and synthesis of α-hydroxytropolones. Med. Chem. Commun. 5, 842–852 (2014).

66. Hadizadeh, I., Peivastegan, B., Hannukkala, A., Wolf, J. M. van der, Nissinen, R. & Pirhonen, M. Biological control of potato soft rot caused by Dickeya solaniand the survival of bacterial antagonists under cold storage conditions. Plant Pathol. 68, 297–311 (2018).

67. Matuszewska, M., Maciąg, T., Rajewska, M., Wierzbicka, A. & Jafra, S. The carbon source-dependent pattern of antimicrobial activity and gene expression in Pseudomonas donghuensis P482. Sci. Rep. 11, 10994 (2021).

68. Agaras, B. C., Scandiani, M., Luque, A., Fernández, L., Farina, F., Carmona, M., Gally, M., Romero, A., Wall, L. & Valverde, C. Quantification of the potential biocontrol and direct plant growth promotion abilities based on multiple biological traits distinguish different groups of Pseudomonas spp. isolates. Biol. Control 90, 173–186 (2015).

69. Agaras, B. C., Iriarte, A. & Valverde, C. F. Genomic insights into the broad antifungal activity, plant-probiotic properties, and their regulation, in Pseudomonas donghuensis strain SVBP6. PLoS ONE 13, e0194088 (2018).

70. Wang, P., Xiao, Y., Gao, D., Long, Y. & Xie, Z. The Gene paaZ of the Phenylacetic Acid (PAA) Catabolic Pathway Branching Point and ech outside the PAA Catabolon Gene Cluster Are Synergistically Involved in the Biosynthesis of the Iron Scavenger 7-Hydroxytropolone in Pseudomonas donghuensis HYS. Int. J. Mol. Sci. 24, 12632 (2023).

71. Darling, A. C. E., Mau, B., Blattner, F. R. & Perna, N. T. Mauve: Multiple Alignment of Conserved Genomic Sequence With Rearrangements. Genome Res. 14, 1394–1403 (2004).

72. Oh, W. T., Jun, J. W., Giri, S. S., Yun, S., Kim, H. J., Kim, S. G., Kim, S. W., Kang, J. W., Han, S. J., Kwon, J., Kim, J. H., Smits, T. H. M. & Park, S. C. Pseudomonas tructae sp. nov., novel species isolated from rainbow trout kidney. Int. J. Syst. Evol. Microbiol. 69, 3851–3856 (2019).

73. Bosi, E., Donati, B., Galardini, M., Brunetti, S., Sagot, M.-F., Lió, P., Crescenzi, P., Fani, R. & Fondi, M. MeDuSa: a multi-draft based scaffolder. Bioinformatics 31, 2443–2451 (2015).

74. Altschul, S. F., Gish, W., Miller, W., Myers, E. W. & Lipman, D. J. Basic local alignment search tool. J. Mol. Biol. 215, 403–410 (1990).

75. Seemann, T. Prokka: rapid prokaryotic genome annotation. Bioinformatics 30, 2068–2069 (2014).

76. Terlouw, B. R., Blin, K., Navarro-Muñoz, J. C., Avalon, N. E., Chevrette, M. G., Egbert, S., Lee, S., Meijer, D., Recchia, M. J. J., Reitz, Z. L., Santen, J. A. van, Selem-Mojica, N., Tørring, T., Zaroubi, L., Alanjary, M., Aleti, G., Aguilar, C., Al-Salihi, S. A. A., Augustijn, H. E., Avelar-Rivas, J. A., Avitia-Domínguez, L. A., Barona-Gómez, F., Bernaldo-Agüero, J., Bielinski, V. A., Biermann, F., Booth, T. J., Bravo, V. J. C., Castelo-Branco, R., Chagas, F. O., Cruz-Morales, P., Du, C., Duncan, K. R., Gavriilidou, A., Gayrard, D., Gutiérrez-García, K., Haslinger, K., Helfrich, E. J. N., Hooft, J. J. J. van der, Jati, A. P., Kalkreuter, E., Kalyvas, N., Kang, K. B., Kautsar, S., Kim, W., Kunjapur, A. M., Li, Y.-X., Lin, G.-M., Loureiro, C., Louwen, J. J. R., Louwen, N. L. L., Lund, G., Parra, J., Philmus, B., Pourmohsenin, B., Pronk, L. J. U., Rego, A., Rex, D. A. B., Robinson, S., Rosas-Becerra, L. R., Roxborough, E. T., Schorn, M. A., Scobie, D. J., Singh, K. S., Sokolova, N., Tang, X., Udwary, D., Vigneshwari, A., Vind, K., Vromans, S. P. J. M., Waschulin, V., Williams, S. E., Winter, J. M., Witte, T. E., Xie, H., Yang, D., Yu, J., Zdouc, M., Zhong, Z., Collemare, J., Linington, R. G., Weber, T. & Medema, M. H. MIBiG 3.0: a community-driven effort to annotate experimentally validated biosynthetic gene clusters. Nucleic Acids Res. 51, D603–D610 (2023).

77. Hmelo, L. R., Borlee, B. R., Almblad, H., Love, M. E., Randall, T. E., Tseng, B. S., Lin, C., Irie, Y., Storek, K. M., Yang, J. J., Siehnel, R. J., Howell, P. L., Singh, P. K., Tolker-Nielsen, T., Parsek, M. R., Schweizer, H. P. & Harrison, J. J. Precision-engineering the Pseudomonas aeruginosa genome with two-step allelic exchange. Nat. Protoc. 10, 1820–1841 (2015).

78. Huang, Y., Niu, B., Gao, Y., Fu, L. & Li, W. CD-HIT Suite: a web server for clustering and comparing biological sequences. Bioinformatics 26, 680–682 (2010).

79. Letunic, I. & Bork, P. Interactive Tree Of Life (iTOL) v5: an online tool for phylogenetic tree display and annotation. Nucleic Acids Res. 49, W293–W296 (2021).

80. Betancor, L., Fernández, M., Weissman, K. J. & Leadlay, P. F. Improved Catalytic Activity of a Purified Multienzyme from a Modular Polyketide Synthase after Coexpression with Streptomyces Chaperonins in Escherichia coli. ChemBioChem 9, 2962–2966 (2008).

