## Supplementary Information for "Understanding the biosynthesis, metabolic regulation, and anti-phytopathogen activity of 3,7-dihydroxytropolone in *Pseudomonas* spp"

### TABLES

**Table S1** Strains used in this study.

| Strain | Genotype / description | Application | Reference |
|---|---|---|---|
| Ps652 | *Pseudomonas* environmental isolate | Environmental biocontrol isolate | 1 |
| Ps652 Δ*hcn* | HCN null mutant (Δ*hcnABC*) | Investigation of Ps652 bioactivity | This work |
| Ps652 Δ*hcn*Δ*pyo* | Pyoverdine null mutant | Investigation of Ps652 bioactivity | This work |
| Ps652ΔNRPS5 | NRPS5 null mutant | Investigation of Ps652 bioactivity | This work |
| Ps652 Δ*hcn* 29C1 | Ps652 Δ*hcn* with transposon insertion mutant *tpoJ* | Investigation of *tpo* gene cluster | This work |
| Ps652 Δ*hcn*Δ*tpoD* | Δ*hcnABC*Δ*tpoD* | Investigation of function of thioesterase TpoD | This work |
| Ps652 Δ*hcn*Δ*tpoE* | Δ*hcnABC*Δ*tpoE* | Investigation of function of dehydrogenase TpoE | This work |
| Ps652 Δ*hcn*Δ*tpoF* | Δ*hcnABC*Δ*tpoF* | Investigation of function of acyl-CoA ligase TpoF | This work |
| Ps652 Δ*hcn*Δ*tpoG* | Δ*hcnABC*Δ*tpoG* | Investigation of function of decarboxylase TpoG | This work |
| Ps652 Δ*hcn*Δ*tpoN* | Δ*hcnABC*Δ*tpoN* | Investigation of function of enoyl-CoA hydratase TpoN | This work |
| Ps652 ΔhcnΔ*tpoD*:: pME6032 | Empty vector control for *tpoD* complementation | Investigation of function of thioesterase TpoD | This work |
| Ps652 ΔhcnΔ*tpoD*:: pME6032-*tpoD* | Δ*hcnABC*Δ*tpoD* complemented with *tpoD* in multicopy expression vector pME6032 | Investigation of function of thioesterase TpoD | This work |
| *Escherichia coli* NEB 5-alpha (Subcloning Efficiency) | *fhuA2 Δ(argF-lacZ)U169 phoA glnV44 Φ80 Δ(lacZ)M15 gyrA96 recA1 relA1 endA1 thi-1 hsdR17* | Plasmid host for molecular cloning | New England Biolabs[2] |
| *Escherichia coli* BL21 (DE3) pL1SL2 | *E. coli* BL21 (DE3) with GroES/GroEL chaperonin system | Protein expression with chaperonins to help protein folding | 3 |
| *Streptomyces scabies* 87-22 | Wild type | Bacterial potato pathogen for assays | 4 |
| *Phytophthora infestans* strain 88069 | Wild type | Oomycete potato pathogen for assays | 5 |

**Table S2** Oligonucleotides used in this study.

| Primer | Sequence 5'-3' | Application |
|---|---|---|
| HCN_UP_F | CAGAGAGGAGCACATTCACG | Amplification of Upstream HCN fragment (forward) |
| HCN_UP_R | CAGTTGATCATGGTCGTTGC | Amplification of upstream HCN fragment (reverse) |
| HCN_DOWN_F | CCTTCCACTGCGATACCCTG | Amplification of downstream HCN fragment (forward) |
| HCN_DOWN_R | CCTGCTTGTTTTCCACAGCC | Amplification of downstream HCN fragment (reverse) |
| tpoD_del_fwd | ACCGAGAGCAGCAGTTCC | PCR screening of Ps652 Δ*tpoD* mutant |
| tpoD_del_rev | ATCAACAGCACTCCTTGAGG | PCR screening of Ps652 Δ*tpoD* mutant |
| tpoD_del_seq | GACCACTTCCTTGCTGTCG | Sequencing of Ps652 Δ*tpoD* mutants |
| tpoF_del_fwd | CGTAGGTGGTCAGTACC | PCR screening of Ps652 Δ*tpoF* mutant |
| tpoF_del_rev | AGGATGCATTGGGCTTCG | PCR screening of Ps652 Δ*tpoF* mutant |
| NRPS_del_fwd | GAACTCGCGATCACTAACC | PCR screening of Ps652 NRPS (BGC6) mutants |
| NRPS_del_rev | CTGGAATCGGATCGAAGTCG | PCR screening of Ps652 NRPS (BGC6) mutants |
| tpoN_del_fwd | CTTACCAGTTTCTGCAGG | PCR screening of Ps652 Δ*tpoN* mutant |
| tpoN_del_rev | GGTGCGATTCATATCG | PCR screening of Ps652 Δ*tpoN* mutant |
| tpoD_comp_ PME6032_F | AAGCCGCCACGGCGATATCGGATCCAT CAACAGCACTCCTTGAGG | Cloning *tpoD* into pME6032 |
| tpoD_comp_ PME6032_R | CCATGGTACCCGGGAGCTCGAATTCTC AGCCTTCCTGGAACG | Cloning *tpoD* into pME6032 |
| tpoG_UP_Fwd | GCTCGGTACCCGGGGATCCTCTAGAAT GGCCGAGCGTGAATGG | Deletion of *tpoG* gene in Ps652 Δ*hcn* |
| tpoG_UP_Rev | GGAATTCGGCATCCGCCATCTGTTTGC CGAGTTCGACTACG | Deletion of *tpoG* gene in Ps652 Δ*hcn* |
| tpoG_DOWN_Fwd | ATGGCGGATGCCGAATTCC | Deletion of *tpoG* gene in Ps652 Δ*hcn* |
| tpoG_DOWN_Rev | GGTGAGAATGGCAAAAGCTCATATGGT CCTGACCACCCTGAGC | Deletion of *tpoG* gene in Ps652 Δ*hcn* |
| tpoG_del_Fwd_2 | GACATCAGCGACGATGACTGG | PCR screening of Ps652 Δ*tpoG* mutants |
| tpoG_del_Rev | TGAGCAAGGAAGCACTGC | PCR screening of Ps652 Δ*tpoG* mutants |
| tpoE_del_Fwd | GTTCACGATGAGCAACAAGG | PCR screening of Ps652 Δ*tpoE* mutants |
| tpoE_del_Rev | GACCACGGTCGGTTTGC | PCR screening of Ps652 Δ*tpoE* mutants |

| Primer | Sequence 5’-3’ | Application |
|---|---|---|
| tpo_comp_UP_F | TCATGCATGAGCTCACTAGTGGATCATC AACAGCACTCCTTGAGG | Amplification of *tpoD* promoter for cloning *tpo* genes into pUC18-mini-Tn7-Gm; PCR screening of complemented mutants |
| tpoE_comp_UP_R | GGAGTCCAGGAGAGTTTCATCGTGAAC GTCCTTTTATTGATTTGTGG | Amplification of tpoD promoter for cloning *tpoE* into pUC18-mini-Tn7-Gm |
| tpoE_comp_DOWN_F | ATGAAACTCTCCTGGACTCC | Cloning *tpoE* into pUC18-mini-Tn7-Gm |
| tpoE_comp_DOWN_R | AGGAATTCCTGCAGCCCGGGGGATCTG GCTCGACCGCTTCAGG | Cloning *tpoE* into pUC18-mini-Tn7-Gm; PCR screening of complemented mutants |
| tpoF_comp_UP_R | GGTATCGATTTGGTTGGCATCGTGAAC GTCCTTTTATTGATTTGTGG | Amplification of tpoD promoter for cloning *tpoF* into pUC18-mini-Tn7-Gm |
| tpoF_comp_DOWN_F | ATGCCAACCAAATCGATACC | Cloning *tpoF* into pUC18-mini-Tn7-Gm |
| tpoF_comp_DOWN_R | AGGAATTCCTGCAGCCCGGGGGATCCT CGTGAGTCCTTTGAGG | Cloning *tpoF* into pUC18-mini-Tn7-Gm; PCR screening of complemented mutants |
| PAA_prom_fwd | CATGCATGAGCTCACTAGTGGATCCTTA GGTTCTTCAGGCCATCC | Cloning *paaA* 5’-UTR into pUC18-mini-Tn7-Gm-lux |
| PAA_prom_rev | GTCATATTTGCCATCCATTTAATGGGGG GTTCTCCAGCCTTTTATGAAGG | Cloning *paaA* 5’-UTR into pUC18-mini-Tn7-Gm-lux |
| PAA_prom_seq_F | GAAGTTCCTATTCCGAAGTTC | Sequencing of pUC18-mini-Tn7-Gm-paa-lux |
| PAA_prom_seq_R | GCAGGTAAACACTATTATCACC | Sequencing of pUC18-mini-Tn7-Gm-paa-lux |
| PAA_prom_quik_F | GGCTGGAGAACCTGGATGGCAAATATG | Site directed mutagenesis of pUC18-mini-Tn7-Gm-paa-lux for optimised distance from RBS to start codon |
| PAA_prom_quik_r | CATATTTGCCATCCAGGTTCTCCAGCC | Site directed mutagenesis of pUC18-mini-Tn7-Gm-paa-lux for optimised distance from RBS to start codon |
| PAA_lux_F | CCACAGCGAATGGATAAGC | Screening of Ps652 clones with integrated pUC18-mini-Tn7-Gm-paa-lux |
| PAA_lux_R | CCAGATAATGGAACATTACCTGC | Screening of Ps652 clones with integrated pUC18-mini-Tn7-G-paa-lux |

**Table S3** Plasmids used in this study.

| Plasmid | Application | Reference |
|---|---|---|
| pTS1 | Suicide vector for in-frame deletions in *Pseudomonas* | [6] |
| pTS1-ΔHCN | In-frame deletion of *hcnABC* gene cluster in Ps652 | This work |
| pTS1-Δ652Pyo | In-frame deletion of pyoverdine encoding NRPS in Ps652 | This work |
| pTS1-ΔNRPS5 | In-frame deletion of L-threonine encoding NRPS in Ps652 | This work |
| pTS1-ΔtpoD | In-frame deletion of *tpoD* in Ps652 *tpo* BGC | This work |
| pTS1-ΔtpoE | In-frame deletion of *tpoE* in Ps652 *tpo* BGC | This work |
| pTS1-ΔtpoF | In-frame deletion of *tpoF* in Ps652 *tpo* BGC | This work |
| pTS1-ΔtpoG | In-frame deletion of *tpoG* in Ps652 *tpo* BGC | This work |
| pTS1-ΔtpoN | In-frame deletion of *tpoN* in Ps652 *tpo* BGC | This work |
| pME6032 | Tetracycline-selective replicative plasmid for IPTG-inducible heterologous expression in *Pseudomonas* | [7] |
| pME6032-tpoD | Complementation of *tpoD* in Ps652Δ*hcn*Δ*tpoD* | This work |
| pUC18-mini-Tn7-Gm | Gentamicin-selective Tn7 integrative vector | [8] |
| pTNS2 | Tn7 transposase expression to facilitate integration of mini-Tn7 vector. | [8] |
| pUC18-mini-Tn7-Gm-*tpoE* | Integrative Tn7 vector for complementation of Ps652 *tpoE* mutants | This work |
| pUC18-mini-Tn7-Gm-*tpoF* | Integrative Tn7 vector for complementation of Ps652 *tpoF* mutants | This work |
| pUC18-mini-Tn7-Gm-lux | Integrative Tn7 vector containing *luxCDABE* genes | [9] |
| pUC18-mini-Tn7-Gm-paa-lux | Integrative Tn7 vector containing *paaA* promoter linked to *luxCDABE* genes | This work |
| pET28b | Gene expression in *E. coli* | Novagen |

**Table S4** Media used in this study

| Medium | Ingredient | Per Litre | Application |
|---|---|---|---|
| Lennox Broth and Lennox Agar (L) | Tryptone | 10 g | Maintenance and growth of *Pseudomonas* strains |
| | Yeast extract | 5 g | |
| | NaCl | 5 g | |
| | D-glucose | 1 g | |
| | Agar (Formedium) | 10 g | |
| Lysogeny Broth (LB) or Lysogeny agar (LB agar) | Bacto-tryptone | 10 g | Maintenance and growth of *E. coli* |
| | Yeast extract | 5 g | |
| | NaCl | 10 g | |
| | Agar (Formedium) | 10 g | |
| Malt Extract-Yeast Extract Maltose Medium (MYM) | Malt extract | 10 g | Cross-streak assays, split-plate assays |
| | Yeast extract | 4 g | |
| | Maltose | 4 g | |
| | Bacteriological agar | 18 g | |
| | Tap water | 1 L | |
| Rye Sucrose Agar (RSA) | Rye grains; pre-germinated, ground using stick blender, heated for 3 hours at 50 °C | 60 g | Biological assays of Ps652 against *P. infestans* |
| | Sucrose | 20 g | |
| Instant Mash Agar (IMA) | Instant mashed potato (Smash, Batchelors) | 20 g | Biological assays of Ps652 fractions against *S. scabies* 87-22; growth of *S. scabies* 87-22 |
| | Lab M #1 agar | 20 g | |
| Modified King's Broth (MKB) | Glycerol | 15 mL | 5 L growth cultures for pyoverdine purification, Transposon mutant metabolomics |
| | $K_2HPO_4$ (anhydrous) | 2.5 g | |
| | Casein hydrolysate | 5 g | |
| | $MgSO_47H_2O$ | 2.5 g | |
| | Adjust to pH 6.7 with HCl | | |
| Modified King's Broth + glucose (MKBG) | Glucose | 15 g | Comparative metabolomics and for 2 L production of tropolones in Ps652 and mutants |
| | $K_2HPO_4$ (anhydrous) | 2.5 g | |
| | Casein hydrolysate | 5 g | |
| | $MgSO_47H_2O$ | 2.5 g | |
| | Adjust to pH 6.7 with HCl | | |
| (CAS) agar | Prepared in accordance with ref. [10] | | Iron binding assays |

**Table S5** Biosynthetic gene clusters in Ps652

| Biosynthetic genes and clusters[1] | antiSMASH detection[2] | GECCO detection[3] | Biosynthetic class | From[4] | To[4] | Predicted product |
|---|---|---|---|---|---|---|
| 1 | No | No | Tropolone | 380,822 | 396,119 | 3,7-dHT |
| 2 | Yes (1.1) | No | N-acetyl glutaminylglutamine amide (NAGGN) | 937,812 | 952,547 | NAGGN |
| 3 | Yes (1.2) | Yes | NRPS | 1,443,088 | 1,528,685 | Pyoverdine[5] |
| 4 | No | No | MddA-like methyltransferase[1] | 1,961,558 | 1,962,307 | Dimethyl sulfide (DMS) |
| 5 | Yes (1.3) | No | RiPP-like (YcaO) | 2,951,482 | 2,963,686 | Unknown peptide |
| 6 | Yes (1.4) | Yes | NRPS | 3,015,864 | 3,055,397 | Unknown peptide |
| 7 | No | Yes | DUF6388[5] | 3,460,872 | 3,467,872 | Unknown (few clear biosynthetic genes[6]) |
| 8 | Yes (1.5) | Yes | NRPS | 3,541,217 | 3,604,665 | Pyoverdine[5] |
| 9 | Yes (1.6) | No | PKS-like (T3PKS or HMG-CoA synthase) | 3,655,571 | 3,696,758 | Unknown |
| 10 | No | Yes | PvcA-like isonitrile synthase[7] | 4,444,442 | 4,452,702 | Unknown |
| 11 | Yes (1.7) | Yes | RiPP-like | 4,493,965 | 4,515,395 | Quinohemoprotein amine dehydrogenase[8] |
| 12 | Yes (1.8) | Yes | T2PKS (aryl polyene) | 5,056,186 | 5,099,787 | Aryl polyene |
| 13 | Yes (1.9) | Yes | NRPS-like | 5,244,418 | 5,287,864 | *mgo*/*pvf*-like operon |
| 14 | No | No | HdtS-like acyl-homoserine lactone synthase[1] | 5,373,110 | 5,373,880 | *N*-acylhomoserine lactone |
| 15 | No | No | HCN | 5,414,905 | 5,417,870 | HCN |
| 16 | No | No | AldA-like indole-3-acetaldehyde dehydrogenase[1] | 5,518,998 | 5,520,491 | Indole-3-acetic acid (IAA) |
| 17 | Yes (1.10) | No | RiPP | 5,884,355 | 5,906,514 | pyrroloquinoline quinone (PQQ) |

1. Genes predicted for the biosynthesis of DMS, *N*-acylhomoserine lactone and IAA are single genes rather than gene clusters. As these molecules can be defined as specialised metabolites, they are listed here.
2. antiSMASH 7.0.0 was used for analysis[11]. Numbers in parentheses refer to the antiSMASH annotation of the BGC. Note that antiSMASH 7.1.0 features rules for the detection of HCN BGCs.
3. https://gecco.embl.de/ v0.9.8 was used[12]. Where relevant, "Yes" indicates where GECCO BGC regions overlap with antiSMASH regions, although the precise boundaries are not identical.
4. antiSMASH or GECCO annotated boundaries are listed. Otherwise, manual annotations of BGC boundaries are based on the location of essential biosynthetic genes.
5. As in other *Pseudomonas* strains, the pyoverdine biosynthetic genes are located in two distinct loci.
6. DUF6388 proteins can be associated with phosphonate BGCs, but no other phosphonate genes are present in this putative cluster.
7. See ref. [13].
8. The quinohemoprotein amine dehydrogenase features a short RiPP peptide with three intrapeptidyl thioether cross-links introduced by a radical SAM protein[14,15].

**Table S6** The tropolone biosynthetic gene cluster in Ps652

| Protein Name | Size (AA) | Locus tag[a] | Protein ID[a] | Pfam domain(s)[b] | Predicted protein activity/domain | Validated function |
|---|---|---|---|---|---|---|
| TpoA | 212 | PS652_00387 | CAK9887586.1 | PF00440<br>PF17918 | TetR family regulator | Positive regulator of biosynthesis in response to low iron[16]. |
| TpoB | 304 | PS652_00386 | CAK9887585.1 | PF03328 | Aldolase/citrate lyase family | |
| TpoC | 246 | PS652_00385 | CAK9887584.1 | PF13561 | NAD(P)H-dependent reductase | |
| TpoD | 137 | PS652_00384 | CAK9887583.1 | PF13279 | Acyl-CoA thioesterase | Essential for biosynthesis; biochemically characterised thioesterase activity. |
| TpoE | 380 | PS652_00383 | CAK9887582.1 | PF02770<br>PF00441<br>PF02771 | Acyl-CoA dehydrogenase / epoxidase | Essential for biosynthesis; biochemically characterised. |
| TpoF | 452 | PS652_00382 | CAK9887581.1 | PF14535<br>PF00501 | Phenylacetate-CoA ligase | Essential for biosynthesis. PAA-CoA production as pathway precursor. |
| TpoG | 563 | PS652_00381 | CAK9887580.1 | PF02776<br>PF00205<br>PF02775 | TPP-dependent decarboxylase | Essential for biosynthesis; phenylacetate biosynthesis. |
| TpoH | 264 | PS652_00380 | CAK9887579.1 | PF13561 | NAD(P)H-dependent reductase | |
| TpoI | 488 | PS652_00379 | CAK9887578.1 | PF02321 | Outer membrane efflux protein | Transposon mutant disrupts biosynthesis[17]. |
| TpoJ | 360 | PS652_00378 | CAK9887577.1 | PF00529 | HlyD-like membrane-fusion protein | |
| TpoK | 519 | PS652_00377 | CAK9887576.1 | PF07690 | Major facilitator superfamily | |
| TpoL | 301 | PS652_00376 | CAK9887575.1 | PF03466<br>PF00126 | LysR family helix-turn-helix regulator | Positive regulator of biosynthesis in response to low iron[16]. |
| TpoM | 150 | PS652_00375 | CAK9887574.1 | PF00582 | Universal stress protein family | |
| TpoN | 258 | PS652_00374 | CAK9887573.1 | PF00378 | Enoyl-CoA hydratase/isomerase | Essential for biosynthesis. Proposed role to intercept PAA catabolon for tropolone biosynthesis. |

[a.] Locus tags and protein IDs are provided for the NCBI/ENA GenBank accession (OZ024668.1) of the *Pseudomonas* sp. Ps652 genome sequence.

[b.] Multiple Pfam domains are listed where they exist in different regions of each protein.

**Table S7** Multistep gradient for semi-preparative scale polar C18 HPLC.

| Retention Time (min) | % $H_2O$ + 0.5% Formic Acid | % MeOH + 0.5% Formic Acid |
|---|---|---|
| 0 | 100 | 0 |
| 5 | 100 | 0 |
| 6 | 85 | 15 |
| 30 | 50 | 50 |
| 35 | 0 | 100 |
| 40 | 0 | 100 |
| 41 | 100 | 0 |
| 46 | 100 | 0 |

### FIGURES

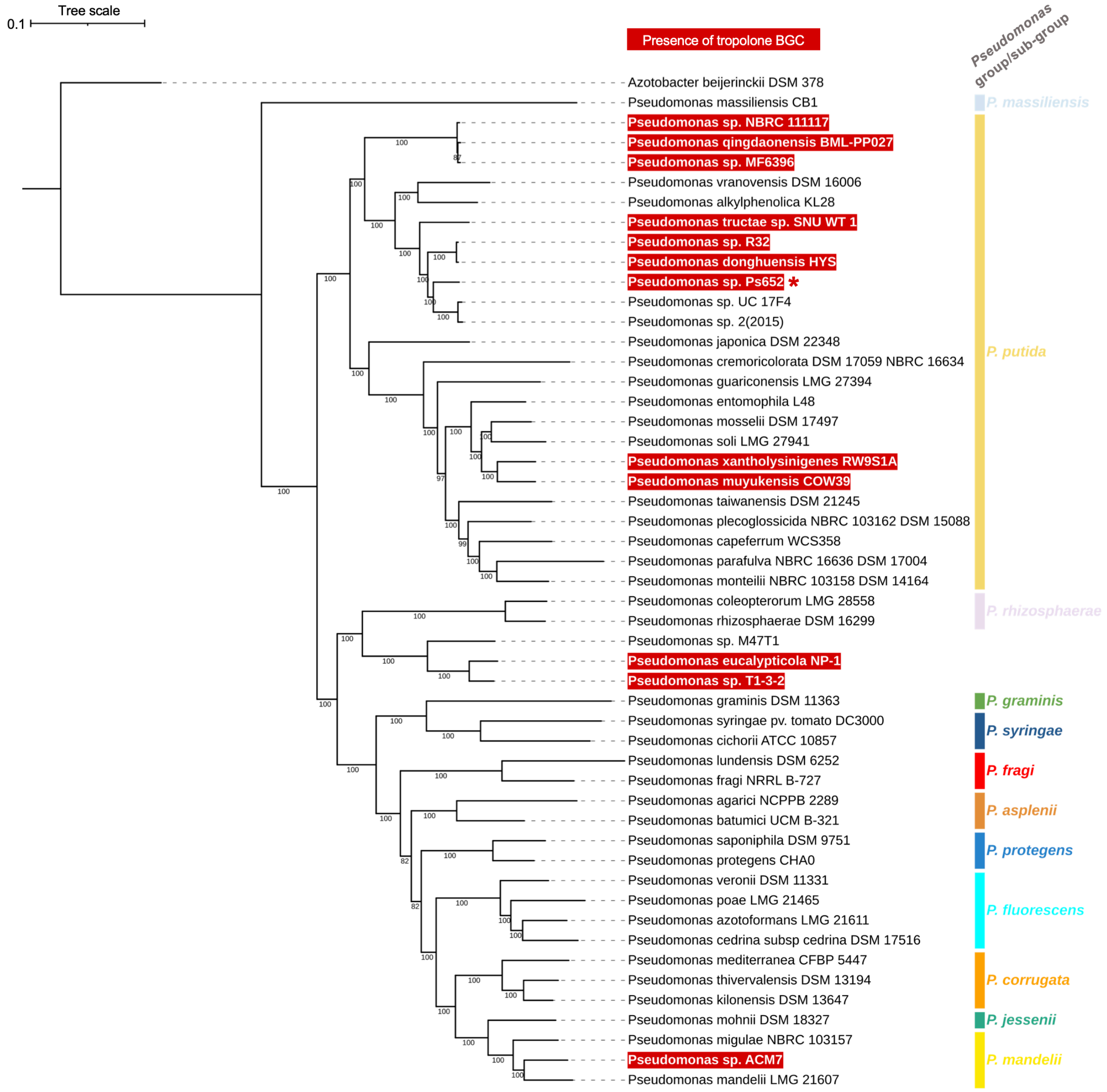

**Figure S1** Phylogenetic analysis of Ps652 versus other *Pseudomonas* strains. Tree generated using AutoMLST[18] and visualised using iTOL[19]. AutoMLST identifies similar species to an input set of genomes to construct a tree featuring a total of 50 representative strains via a concatenated alignment of 83 conserved genes. Strains highlighted with a white text on a red background have a tropolone BGC, while *Pseudomonas* groups/subgroups described in Girard *et al.*[20] are highlighted by coloured strips. Girard *et al.* proposed that the strains within the Ps652 clade (including *P. donghuensis*, *P. tructae* and *P. vranovensis*) belong to the *P. vranovensis* sub-group. The clade containing *Pseudomonas eucalypticola* is not closely related enough (via average nucleotide identity) to *P. rhizosphaerae* to place these strains in the *P. rhizosphaerae* group so these strains are not associated with a specific group. Ps652 is highlighted with an asterisk.

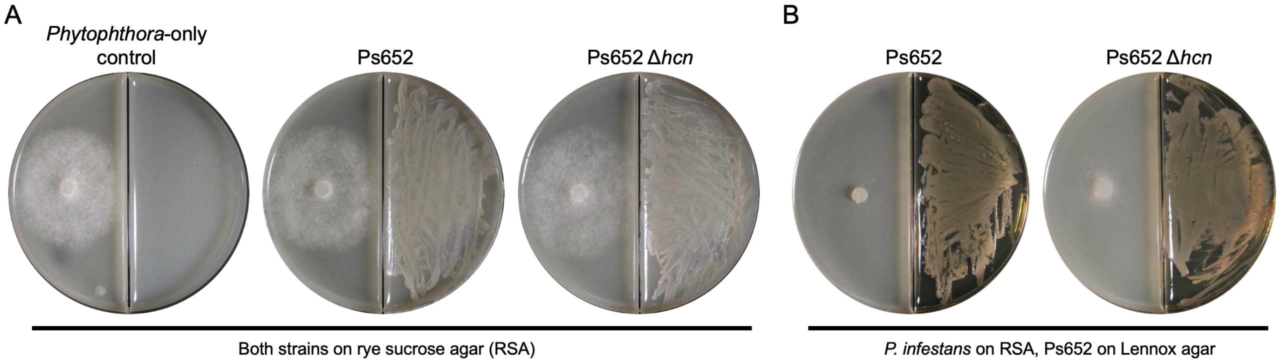

**Figure S2** Split-plate assays for Ps652 and Ps652 Δ*hcn* versus *P. infestans*. **A.** Both pathogen and pseudomonad are plated on RSA. Little inhibition is observed, and no difference in inhibition is evident between the wild type Ps652 and the cyanide mutant Ps652 Δ*hcn*. **B.** Split-plate assays for Ps652 and Ps652 Δ*hcn* versus *P. infestans* with Ps652 strains plated on Lennox agar. Strong volatile production on Lennox agar provides complete inhibition of *P. infestans* by wild type Ps652. Some activity is lost in the cyanide mutant Ps652 Δ*hcn*, but significant inhibition of P. infestans remains, indicating at least one further volatile produced on Lennox agar. All images are representative of three biological replicates.

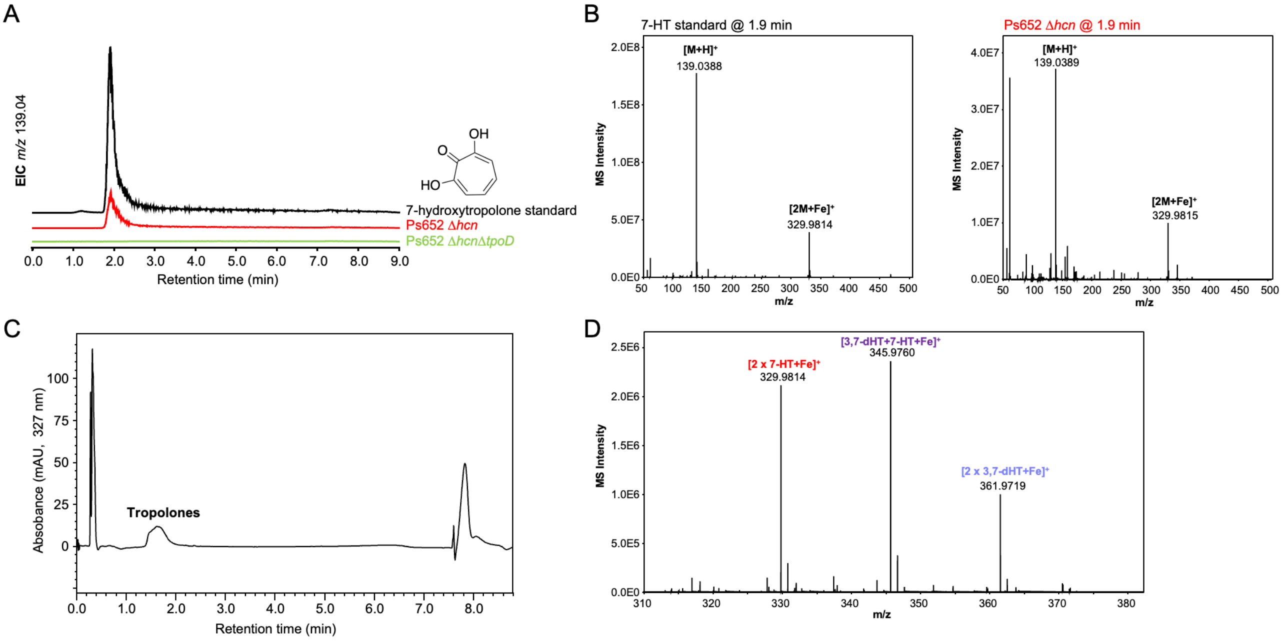

**Figure S3** Detection of tropolones in Ps652. **A.** Production of 7-hydroxytropolone in Ps652 Δ*hcn* in comparison to a synthetic standard provided by Prof. Ryan Murelli (Brooklyn College, The City University of New York). Production is lost in the Δ*tpoD* mutant. EIC = extracted ion chromatogram. B. Comparison of MS spectra for standard 7-HT and the molecule produced in Ps652. C. UV chromatogram for the data shown in Figure 3B of the paper. D. MS spectrum showing the three $[2M+Fe]^+$ adducts of 7-HT and 3,7-dHT.

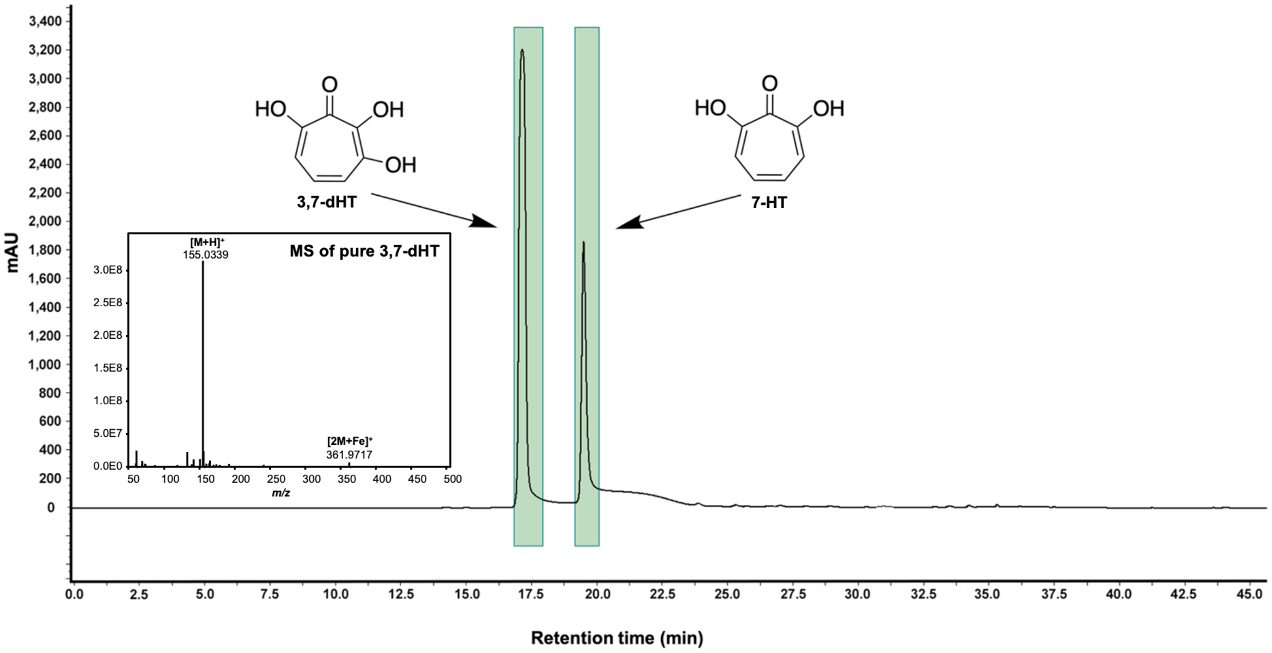

**Figure S4** Separation of 7-HT and 3,7-dHT using semi-preparative HPLC. Trace shows UV absorbance at 327 nm. Inset, MS spectrum for purified 3,7-dHT.

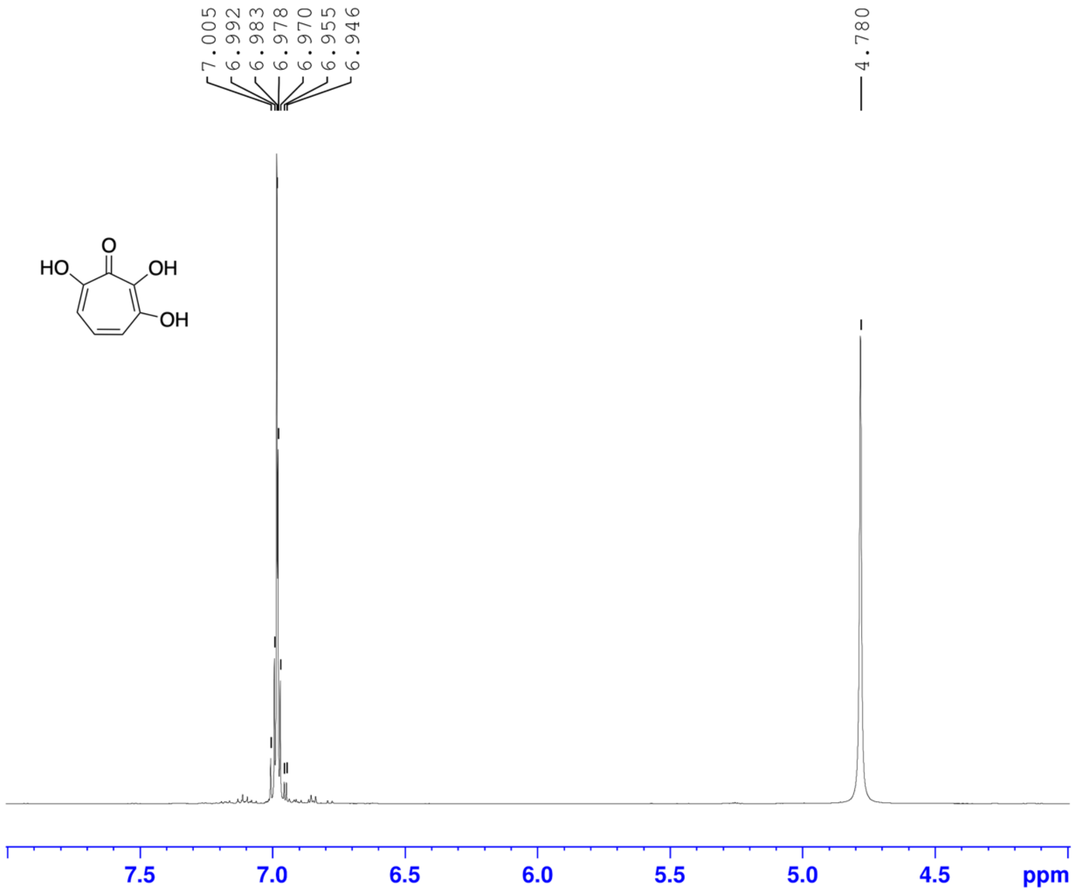

**Figure S5** [1]H NMR spectrum (600 MHz, $CD_3OD$, 298 K) of 3,7-dHT. The multiplet at 6.98 ppm was identified as the 3 CH protons, which matches with to a previously published spectrum for this compound[21], where a singlet was observed due to the lower resolution instrument used for that study. The broad singlet peak at 4.78 ppm is the solvent peak.

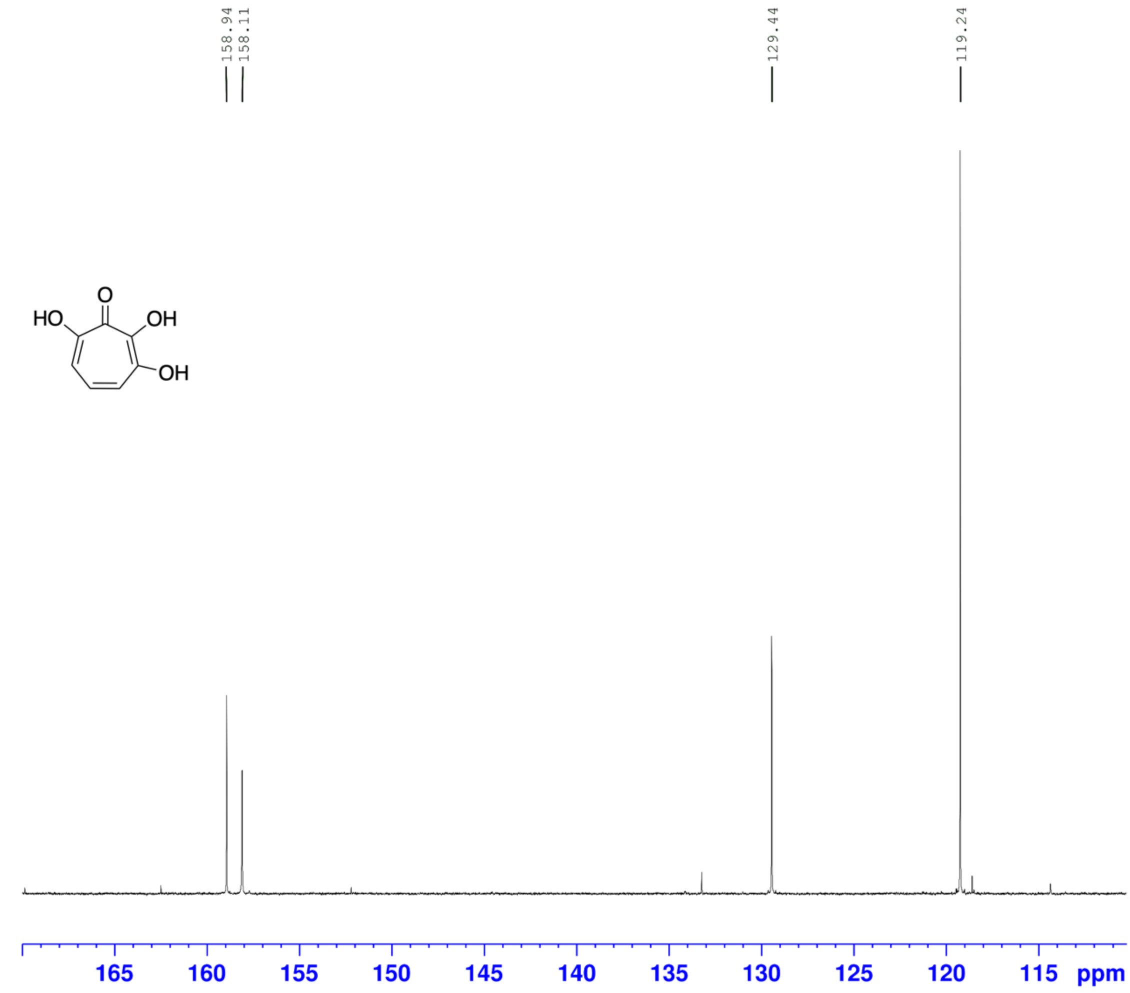

**Figure S6** $^{13}C$ NMR spectrum (150 MHz, $CD_3OD$, 298 K) of 3,7-dHT.

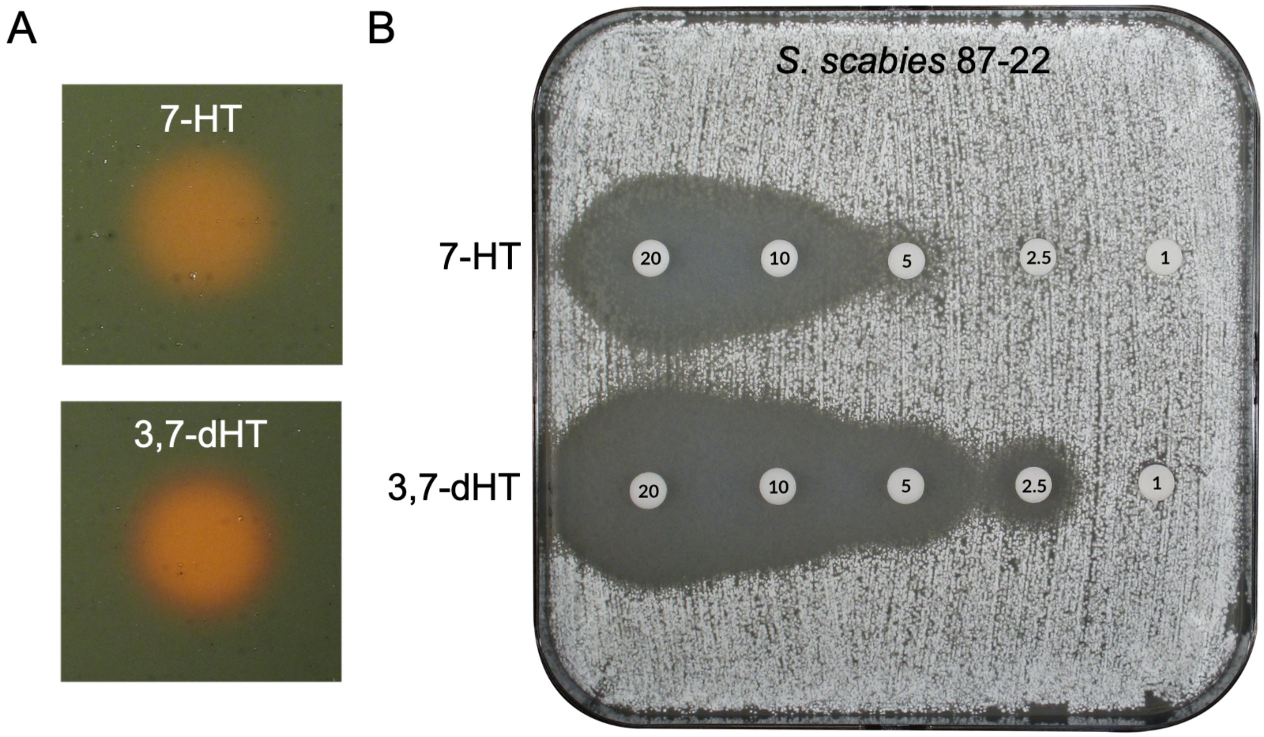

**Figure S7** Comparison of antibacterial and iron binding activities of 7-HT and 3,7-dHT. **A.** Binding of 7-HT and 3,7-dHT to iron on chrome azurol S (CAS) agar plates. 5 µg each of 7-HT and 3,7-dHT were added to the surface of a CAS agar plate in 5 µL methanol. Both molecules displayed the ability to bind iron as indicated by the blue to orange colour change. B. Minimum inhibitory concentration (MIC) assay of 7-HT and 3,7-dHT with *S. scabies* 87-22. Numbers refer to micrograms of molecule in each paper disc.

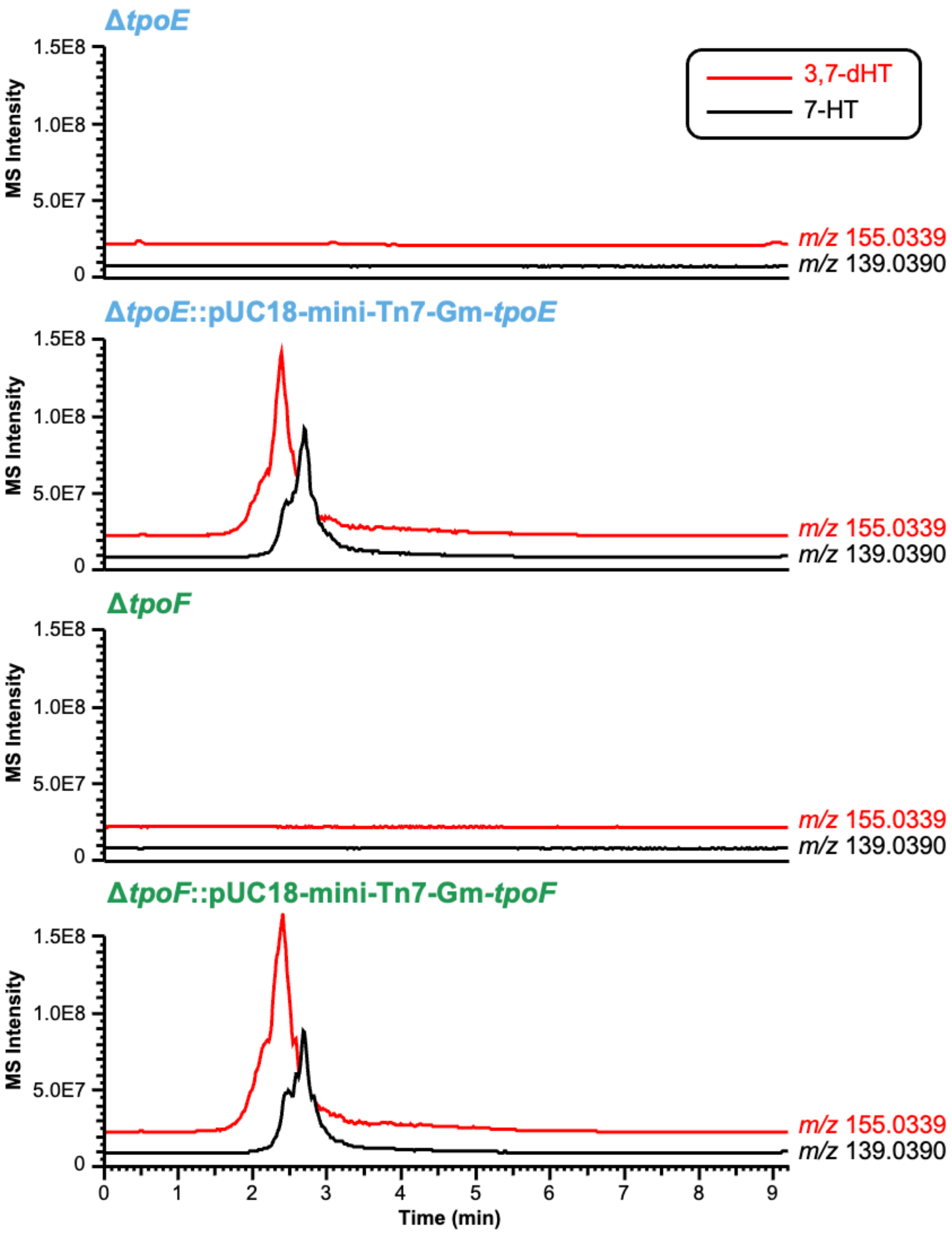

**Figure S8** LC-MS chromatograms of Δ*tpoE* and Δ*tpoF* alongside strains complemented using pME6032-based plasmids. Each mass is searched within a ± *m/z* 0.007 window.

A

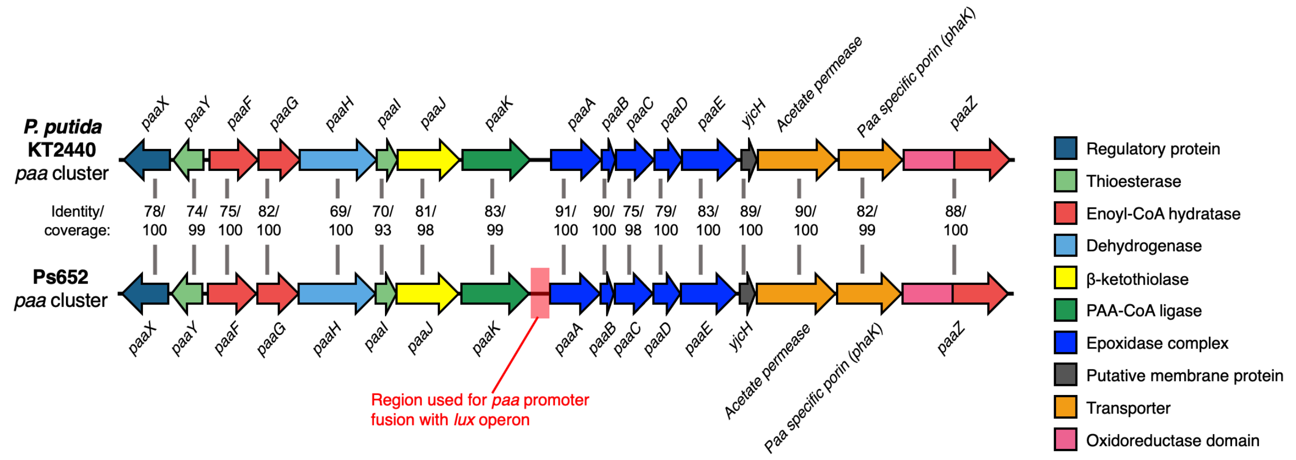

B

TTAGGTTCTTCAGGCCATCCCTGGGCCATGGGCCCTGGGATGGATGTTGAGGCTGCCAGCTTATCCATTCGTT
GTGGCGTGGCTGCCGACCCCTTCCGCCCTTACGGCGGCCTACTTTTCTCTTGGGAGGAAGTAGGCAAAACCGC
TTGCTCCTGTATACGGCCCTACGCTGCGCTTCGGGTCCCTTCGCTACGGCACCCCGCAAGAGCGCCACAGCGA
ATGGATAAGCGCTCCGTTACGCCCGAACAAAGCGATACACAAATCAGTTCTGATACACATTTTTGTGTTTGAT
ATTTTGCCCTGTATCACATATAACTGTCTCCAGCCTCAAAGCCCCCTTCATAAAAGGCTGGAGAACCCC

**Figure S9** Overview of the PAA catabolon. A. Comparison of the PAA catabolon genes in *Pseudomonas putida* KT2440 and Ps652. The promoter region used for the *lux* reporter assays is highlighted in red. Where relevent, the colour coding for general protein activities is consistent with the *tpo* BGC. B. The sequence upstream of *paaA* in Ps652 used in the promoter fusion.

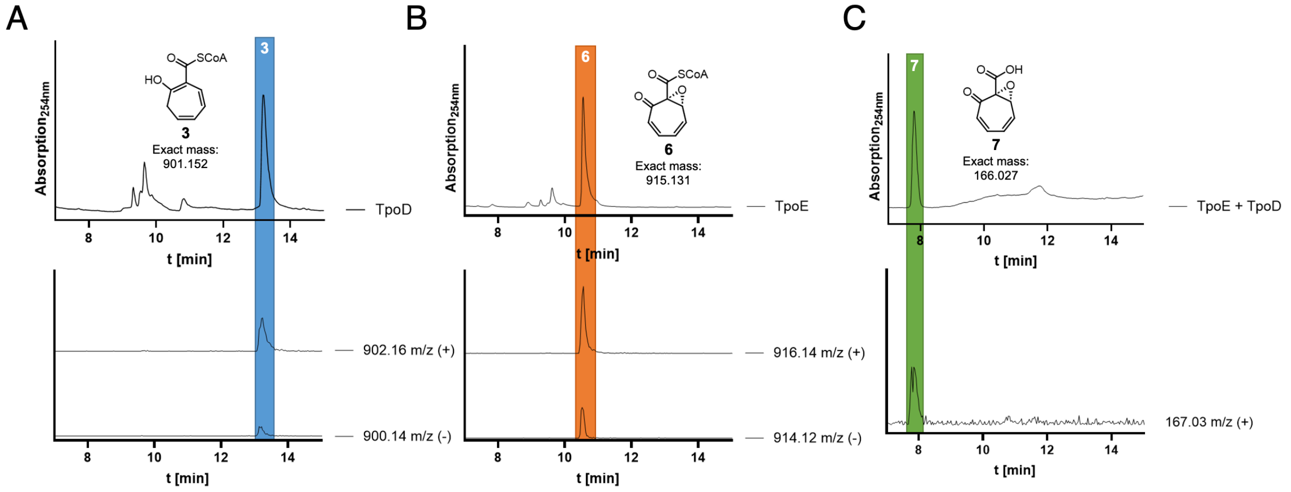

**Figure S10** LC-MS analysis of TpoD, TpoE and TpoE + TpoD products from biochemical reactions. A. Positive and negative EICs of compound **3**. B. Positive and negative EICs for the TpoE product resulting from the conversion of **3**. C. Positive EIC for compound **7** generated by incubation with TpoE and TpoD (only positive mode shown due to poor ionization in negative mode).

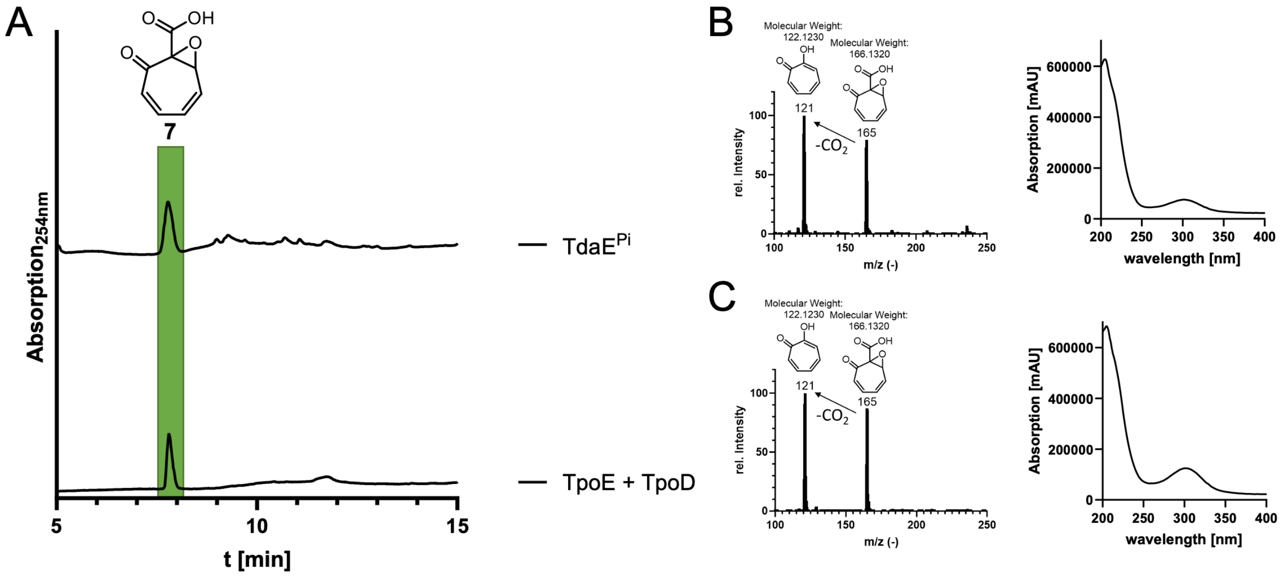

**Figure S11** Comparison between compound **7** produced by TdaE$^{Pi}$ and TpoE + TpoD. TdaE$^{Pi}$ has previously been shown to catalyse the conversion of **3** into **7**.[22] A. HPLC analysis at 254 nm of compound **7** (highlighted in green) at 254 nm produced by TpoE + TpoD (bottom trace) or TdaE$^{Pi}$ (top trace). MS pattern with characteristic internal fragmentation (resulting from decarboxylation) and UV-visible spectrum of compound **7** produced by (**B**) TdaE$^{Pi}$ or (**C**) TpoE + TpoD.

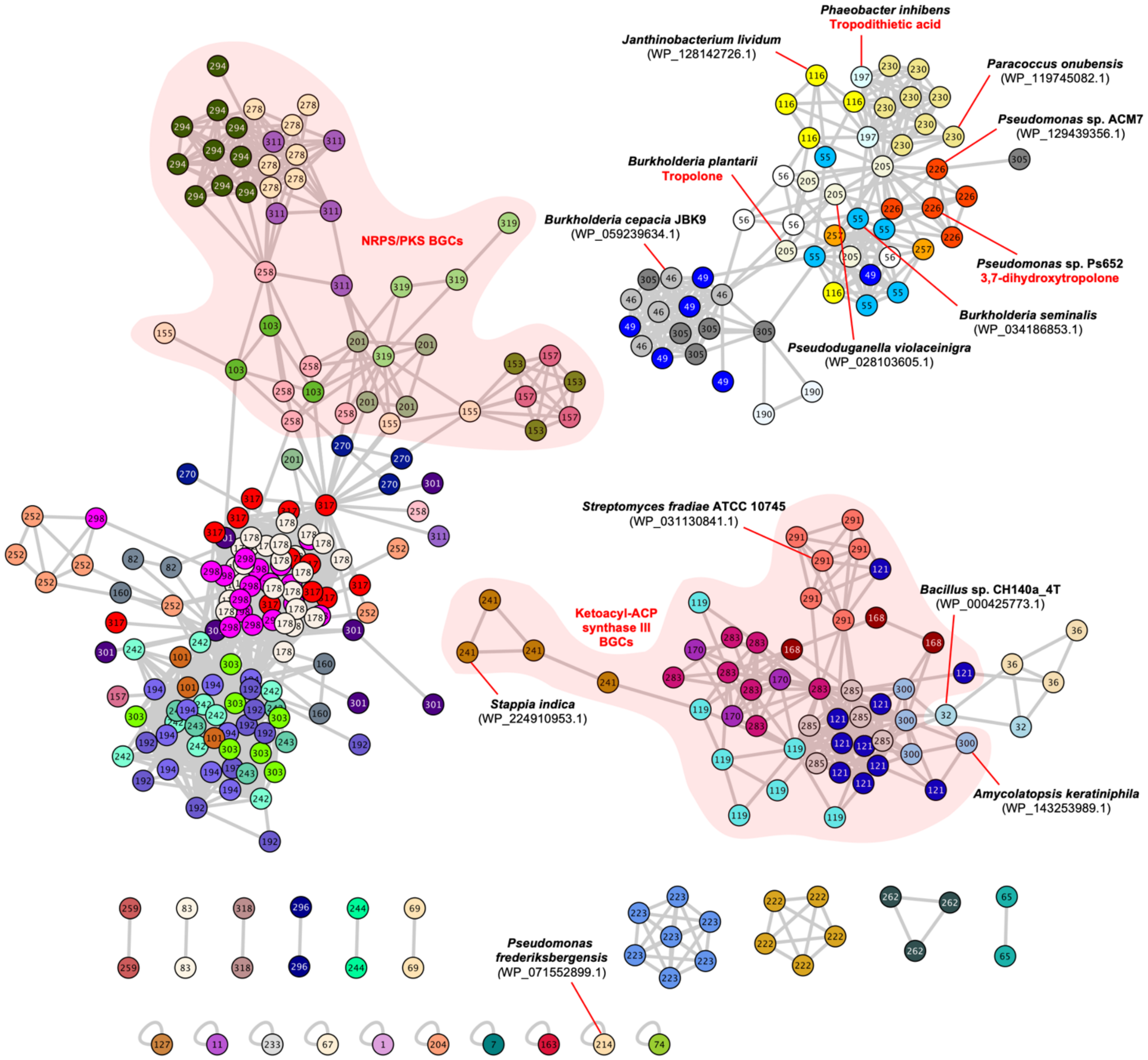

**Figure S12** BiG-SCAPE[23] networking of BGCs retrieved from TpoE/TdaE-based genome mining. BGCs are labelled according to their family within the "Others" network with a 0.75 cut-off and are colour-coded to distinguish between families. The BGCs shown in Figure 6 are labelled, along with sets of BGCs that encode NRPS and/or PKS proteins alongside the flavoproteins (Figure S11). The BiG-SCAPE networks are visualised using Cytoscape.

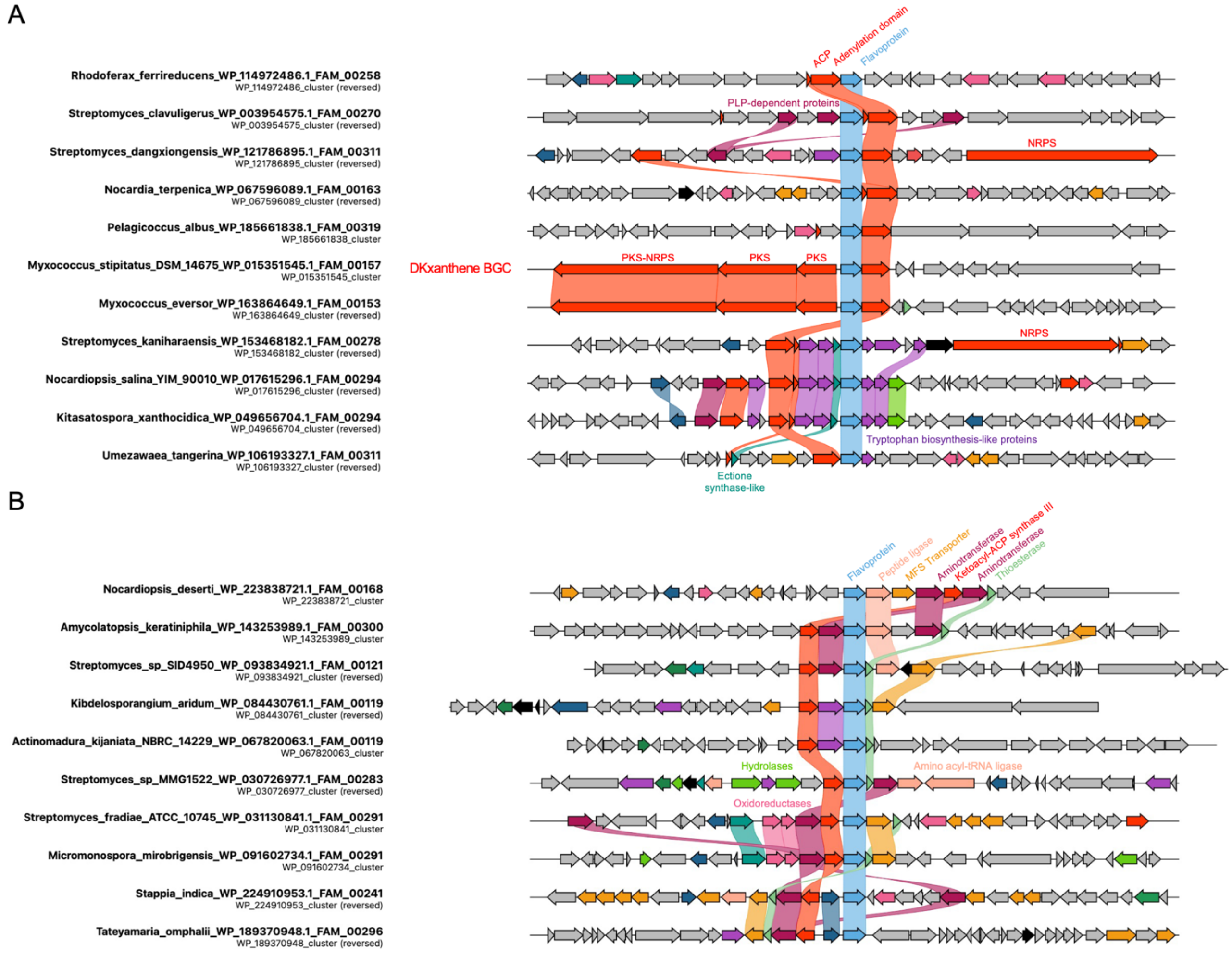

**Figure S13** Examples of flavoprotein-associated BGCs that encode NRPS and/or PKS proteins. Family numbers, flavoprotein accession and species are listed with each BGC. Note that only genes with homology from the clinker analysis[24] (identity threshold of 30%) are colour-coded (see Supplementary File 2), so some BGCs do feature further assembly line genes, and some BGCs extend beyond the regions visible in the figure due to the criteria for BGC retrieval (35 kb regions centred on the flavoprotein gene). A. Example BGCs featuring NRPS or hybrid PKS-NRPS proteins. The BGC in *Myxococcus stipitatus* DSM 14675 is responsible for the production of DKxanthene[25] where it is proposed that the flavoprotein functions as a prolyl dehydrogenase[26]. B. Example BGCs that encode a conserved ketoacyl-ACP synthase III protein alongside multiple other conserved biosynthetic proteins, including a thioesterase.

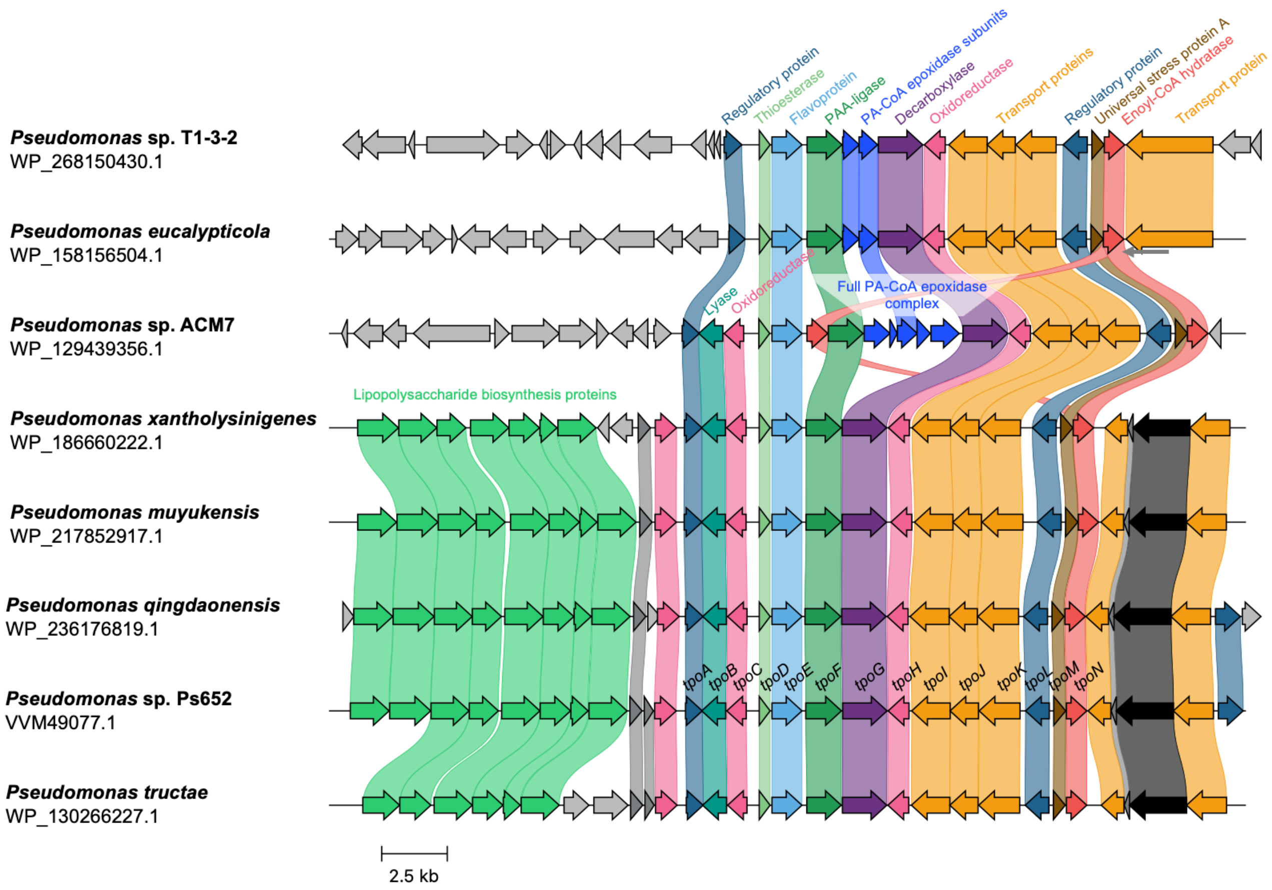

**Figure S14** Comparison of tropolone BGCs in the genus *Pseudomonas*. These BGCs represent the diversity of BGCs with TpoE homologues with >50% identity. A 99% identity filter was used to reduce the redundancy of the dataset without limiting the diversity of output. A subset of BGCs encode some or all of the PA-CoA epoxidase subunits involved in the PAA catabolon, while *Pseudomonas* sp. ACM7 also has an extra copy of an enoyl-CoA hydratase gene. The *tpo* genes are listed for the Ps652 BGC and the accessions for the TpoE homologues are listed under each species name.

### REFERENCES

1. Pacheco-Moreno, A., Stefanato, F. L., Ford, J. J., Trippel, C., Uszkoreit, S., Ferrafiat, L., Grenga, L., Dickens, R., Kelly, N., Kingdon, A. D., Ambrosetti, L., Nepogodiev, S. A., Findlay, K. C., Cheema, J., Trick, M., Chandra, G., Tomalin, G., Malone, J. G. & Truman, A. W. Pan-genome analysis identifies intersecting roles for Pseudomonas specialized metabolites in potato pathogen inhibition. *eLife* **10,** e71900 (2021).

2. Anton, B. P. & Raleigh, E. A. Complete Genome Sequence of NEB 5-alpha, a Derivative of Escherichia coli K-12 DH5α. *Genome Announc.* **4,** e01245-16 (2016).

3. Betancor, L., Fernández, M., Weissman, K. J. & Leadlay, P. F. Improved Catalytic Activity of a Purified Multienzyme from a Modular Polyketide Synthase after Coexpression with Streptomyces Chaperonins in Escherichia coli. *ChemBioChem* **9,** 2962–2966 (2008).

4. Loria, R., Bukhalid, R. A., Creath, R. A., Leiner, R. H., Olivier, M. & Steffens, J. C. Differential production of thaxtomins by pathogenic Streptomyces species in vitro. *Phytopathology* **85,** 537–541 (1995).

5. Kamoun, S., West, P. van, Vleeshouwers, V. G. A. A., Groot, K. E. de & Govers, F. Resistance of Nicotiana benthamiana to Phytophthora infestans Is Mediated by the Recognition of the Elicitor Protein INF1. *Plant Cell* **10,** 1413 (1998).

6. Scott, T. A., Heine, D., Qin, Z. & Wilkinson, B. An L-threonine transaldolase is required for L-threo-β-hydroxy-α-amino acid assembly during obafluorin biosynthesis. *Nat. Commun.* **8,** 15935 (2017).

7. Heeb, S., Itoh, Y., Nishijyo, T., Schnider, U., Keel, C., Wade, J., Walsh, U., O'Gara, F. & Haas, D. Small, stable shuttle vectors based on the minimal pVS1 replicon for use in gram-negative, plant-associated bacteria. *Mol. Plant-Microbe Interact.* **13,** 232–237 (2000).

8. Choi, K.-H., Gaynor, J. B., White, K. G., Lopez, C., Bosio, C. M., Karkhoff-Schweizer, R. R. & Schweizer, H. P. A Tn7-based broad-range bacterial cloning and expression system. *Nat. Meth.* **2,** 443–448 (2005).

9. Glassing, A. & Lewis, T. A. An improved Tn7-lux reporter for broad host range, chromosomally-integrated promoter fusions in Gram-negative bacteria. *J. Microbiol. Methods* **118,** 75–77 (2015).

10. Louden, B. C., Haarmann, D. & Lynne, A. M. Use of Blue Agar CAS Assay for Siderophore Detection. *J. Microbiol. Biol. Educ.* **12,** 51–53 (2011).

11. Blin, K., Shaw, S., Augustijn, H. E., Reitz, Z. L., Biermann, F., Alanjary, M., Fetter, A., Terlouw, B. R., Metcalf, W. W., Helfrich, E. J. N., van Wezel, G. P., Medema, M. H. & Weber, T. antiSMASH 7.0: new and improved predictions for detection, regulation, chemical structures and visualisation. *Nucleic Acids Res.* **51,** W46–W50 (2023).

12. Carroll, L. M., Larralde, M., Fleck, J. S., Ponnudurai, R., Milanese, A., Cappio, E. & Zeller, G. Accurate de novo identification of biosynthetic gene clusters with GECCO. *BioRxiv* 2021.05.03.442509 (2021). doi:10.1101/2021.05.03.442509

13. Clarke-Pearson, M. F. & Brady, S. F. Paerucumarin, a New Metabolite Produced by the pvc Gene Cluster from Pseudomonas aeruginosa. *J. Bacteriol.* **190,** 6927–6930 (2008).

14. Vandenberghe, I., Kim, J.-K., Devreese, B., Hacisalihoglu, A., Iwabuki, H., Okajima, T., Kuroda, S., Adachi, O., Jongejan, J. A., Duine, J. A., Tanizawa, K. & Beeumen, J. V. The Covalent Structure of the Small Subunit from Pseudomonas putida Amine Dehydrogenase Reveals the Presence of Three Novel Types of Internal Cross-linkages, All Involving Cysteine in a Thioether Bond. *J. Biol. Chem.* **276,** 42923–42931 (2001).

15. Nakai, T., Deguchi, T., Frébort, I., Tanizawa, K. & Okajima, T. Identification of Genes Essential for the Biogenesis of Quinohemoprotein Amine Dehydrogenase. *Biochemistry* **53,** 895–907 (2014).

16. Chen, M., Wang, P. & Xie, Z. A Complex Mechanism Involving LysR and TetR/AcrR That Regulates Iron Scavenger Biosynthesis in Pseudomonas donghuensis HYS. *J. Bacteriol.* **200,** 592 (2018).

17. Moffat, A. D., Elliston, A., Patron, N. J., Truman, A. W. & Lopez, J. A. C. A biofoundry workflow for the identification of genetic determinants of microbial growth inhibition. *Synth. Biol.* **6,** ysab004 (2021).

18. Alanjary, M., Steinke, K. & Ziemert, N. AutoMLST: an automated web server for generating multi-locus species trees highlighting natural product potential. *Nucleic Acids Res.* **47,** W276–W282 (2019).

19. Letunic, I. & Bork, P. Interactive Tree Of Life (iTOL) v5: an online tool for phylogenetic tree display and annotation. *Nucleic Acids Res.* **49,** W293–W296 (2021).

20. Girard, L., Lood, C., Höfte, M., Vandamme, P., Rokni-Zadeh, H., Noort, V. van, Lavigne, R. & Mot, R. D. The Ever-Expanding Pseudomonas Genus: Description of 43 New Species and Partition of the Pseudomonas putida Group. *Microorganisms* **9,** 1766 (2021).

21. Takeshita, H., Mori, A., Kusaba, T. & Watanabe, H. Preparation of Polyacetoxytropones and Polyhydroxytropolones by Acetolysis and Hydrolysis of Halotroponoids by Acetyl Trifluoroacetate with Exhaustive Displacement of Halogens on the Tropone Ring. Predominant Formation of Reductive Acetolysates from Fully-Substituted Tropones. *Bull. Chem. Soc. Jpn.* **60,** 4325–4333 (1987).

22. Duan, Y., Toplak, M., Hou, A., Brock, N. L., Dickschat, J. S. & Teufel, R. A Flavoprotein Dioxygenase Steers Bacterial Tropone Biosynthesis via Coenzyme A-Ester Oxygenolysis and Ring Epoxidation. *J. Am. Chem. Soc.* **143,** 10413–10421 (2021).

23. Navarro-Muñoz, J. C., Selem-Mojica, N., Mullowney, M. W., Kautsar, S. A., Tryon, J. H., Parkinson, E. I., Santos, E. L. C. de L., Yeong, M., Cruz-Morales, P., Abubucker, S., Roeters, A., Lokhorst, W., Fernandez-Guerra, A., Cappelini, L. T. D., Goering, A. W., Thomson, R. J., Metcalf, W. W., Kelleher, N. L., Barona-Gómez, F. & Medema, M. H. A computational framework to explore large-scale biosynthetic diversity. *Nat. Chem. Biol.* **16,** 60–68 (2020).

24. Gilchrist, C. L. M. & Chooi, Y.-H. clinker & clustermap.js: automatic generation of gene cluster comparison figures. *Bioinformatics* **37,** 2473–2475 (2021).

25. Hyun, H., Lee, S., Lee, J. S. & Cho, K. Genetic and functional analysis of the DKxanthene biosynthetic gene cluster from Myxococcusstipitatus DSM 14675. *J. Microbiol. Biotechnol.* **28,** 1068–1077 (2018).

26. Meiser, P., Weissman, K. J., Bode, H. B., Krug, D., Dickschat, J. S., Sandmann, A. & Müller, R. DKxanthene Biosynthesis—Understanding the Basis for Diversity-Oriented Synthesis in Myxobacterial Secondary Metabolism. *Chem. Biol.* **15,** 771–781 (2008).
