## Supplementary figures and images for "Understanding the biosynthesis, metabolic regulation, and anti-phytopathogen activity of 3,7-dihydroxytropolone in *Pseudomonas* spp"

### Supplementary File 2

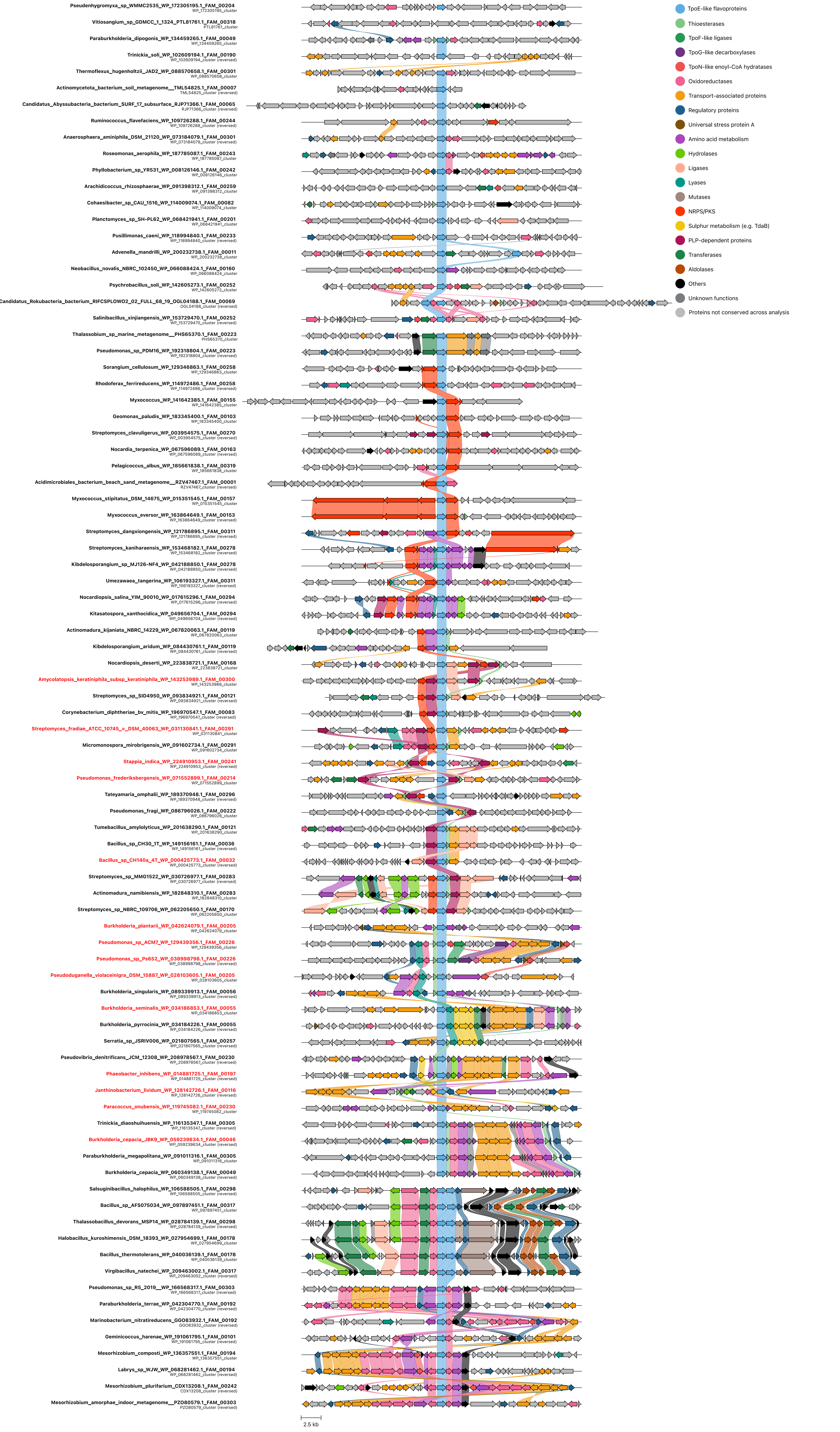
